# KIPEs3: Automatic annotation of biosynthesis pathways

**DOI:** 10.1101/2022.06.30.498365

**Authors:** Andreas Rempel, Nancy Choudhary, Boas Pucker

## Abstract

Flavonoids and carotenoids are pigments involved in stress mitigation and numerous other processes. Both pigment classes can contribute to flower and fruit coloration. Flavonoid aglycones and carotenoids are produced by a pathway that is largely conserved across land plants. Glycosylations, acylations, and methylations of the flavonoid aglycones can be species-specific and lead to a plethora of biochemically diverse flavonoids. We previously developed KIPEs for the automatic annotation of biosynthesis pathways and presented an application on the flavonoid aglycone biosynthesis.

KIPEs3 is an improved version with additional features and the potential to identify not just the core biosynthesis players, but also candidates involved in the decoration steps and in the transport of flavonoids. Functionality of KIPEs3 is demonstrated through the analysis of the flavonoid biosynthesis in *Arabidopsis thaliana* Nd-1, *Capsella grandiflora*, and *Dioscorea dumetorum*. We demonstrate the applicability of KIPEs to other pathways by adding the carotenoid biosynthesis to the repertoire. As a technical proof of concept, the carotenoid biosynthesis was analyzed in the same species and *Daucus carota*. KIPEs3 is available as an online service to enable access without prior bioinformatics experience.

KIPEs3 facilitates the automatic annotation and analysis of biosynthesis pathways with a consistent and high quality in a large number of plant species. Numerous genome sequencing projects are generating a huge amount of data sets that can be analyzed to identify evolutionary patterns and promising candidate genes for biotechnological and breeding applications.

## Background

Carotenoids and flavonoids are two important classes of plant pigments. Colorful flowers are often pigmented by one or both of these pigments. Flavonoids (anthocyanins) are known for blue, purple, red, and orange colors, while carotenoids can provide yellow, orange, and red pigmentation. Biological roles of carotenoids include photoprotection [1, 2], light capture [3], coloration of plant structures [4–6], and hormone precursors [7]. Therefore, carotenoids are considered part of the primary metabolism. In contrast, flavonoids are specialized plant metabolites that contribute to stress resilience and play important roles in the reproduction of many plants [8–10]. The biosynthesis of flavonoid aglycones [11, 12], the decoration with sugar and acyl groups [13–15], and the intracellular transport of flavonoids [16–18] have been reviewed elsewhere. The core of the flavonoid biosynthesis pathway is well conserved across land plant species, whereas the decorating reactions are often species-specific. Glycosylation and acylation modify the properties of flavonoids, but specific properties depend on the added sugar moiety or acyl group. Glycosylation can enhance the anthocyanin stability [19–21] and be required for intracellular flavonoid transport [17, 18]. Acylation can also boost the stability of anthocyanins [22]. Applications of anthocyanins as food colorants rely on a high stability.

Biotechnological applications involving flavonoids are generally facilitated by knowledge about the genes involved in their biosynthesis. For example, such insights are crucial for the production of flavonoids in heterologous systems [23]. The flavonoid biosynthesis pathway is well studied in several model organisms and crops like *Arabidopsis thaliana* [18, 20], *Petunia hybrida* [11, 24, 25], *Antirrhinum majus* [26, 27], *Vitis vinifera* [28–30], and *Zea mays* [20, 31]. An ongoing challenge is the efficient transfer of this knowledge to other plant species. In fact, the functional annotation transfer challenge goes far beyond knowledge about the flavonoid biosynthesis. Plant genome sequences are becoming available with an increasing pace [32, 33], but the functional annotation is lacking behind. Automatic approaches for biological interpretation of these data sets are crucial to exploit the full value of genomic resources. There are several comprehensive databases collecting and sharing knowledge about plant metabolism, e.g. The Plant Metabolic Network [34], Plant Reactome [35], and MetaCyc [36]. Tools like InterProScan5 [37], Blast2GO [38], KEGG Automatic Annotation Server [39], and Mercator4 [40] allow the assignment of certain annotation terms to input sequences. While these tools are able to annotate genome-scale data sets, they are less specific with respect to the assigned annotation. Researchers looking for a specific pathway could benefit from tools focusing on their pathway of interest.

Here, we present an updated approach for the automatic annotation of the flavonoid biosynthesis genes in any land plant species of interest based on prior knowledge about the pathway. We also demonstrate that this approach can be applied to other pathways by investigating the carotenoid biosynthesis genes in multiple land plant species. A web server makes this automatic annotation workflow available to the research community.

## Materials and Methods

### Validation of KIPEs3 results through manual inspection

A validation of KIPEs3 was based on *Arabidopsis thaliana* Nd-1 v1.0 [41], *Capsella grandiflora* v1.1 [42], *Daucus carota* v2.0 [43], and *Dioscorea dumetorum* v1.0 [44]. Identified candidates were manually inspected in a phylogenetic tree constructed by FastTree2 [45] based on alignments generated with MAFFT v7 [46].

### Run time analysis of KIPEs3

The average run time of KIPEs3 per data set was assessed using a previously defined set of 121 plant species [47]. All these sequence files were retrieved from Phytozome [48]. The analysis was performed on coding sequences to represent transcriptome assemblies. Performance of KIPEs3 with FastTree v2.1.10 [45] was analyzed using default parameters. Run time and the number of sequences in the analyzed data sets were documented. Run times were measured for analyses with the construction of phylogenetic trees for all final candidate genes.

### Comparison of KIPEs3 predictions against annotation databases

The accuracy of KIPEs3 predictions was compared against MetaCyc [49], Plant Reactome [35], and Plant Metabolic Network (PMN) [34]. *FLS* was selected as a representative gene of the flavonoid biosynthesis and *LYC-b* was selected as a representative gene of the carotenoid biosynthesis. The compared species were determined by overlap between the databases with respect to the covered species. The usage of the same genome sequence and annotation for a species across different databases was important. *FLS* annotations in *Arabidopsis thaliana, Cicer arietinum*, and *Citrus sinensis* were compared. The *LYC-b* annotations in *A. thaliana, Solanum lycopersicum*, and *Daucus carota* were compared. The candidates were searched using the Enzyme Commission numbers, EC 1.14.20.6 for *FLS* and EC 5.5.1.19 for *LYC-b* in Metacyc and PMN. In Plant Reactome, the genes were searched using the terms “flavonol synthase” and “lycopene beta cyclase” in the search bar. KIPEs3 v0.35 was run as a command-line tool providing flavonoid biosynthesis baits, residues, and the predicted peptide sequences of the species to identify *FLS* candidates in each species. The *LYC-b* candidates were annotated in another run based on the carotenoid biosynthesis bait sequences, residues, and the predicted peptide sequences of the respective species. The *FLS* and *LYC-b* candidates listed in the three databases and identified by KIPEs3 were aligned using MAFFT v7 [46] to discriminate functional proteins and the products of pseudogenes by identifying *FLS* and *LYC-b* specific motifs and residues, respectively. A phylogenetic tree of the *LYC-b* candidates was generated using FastTree v2.1 [45].

### Gene expression analysis

Gene expression was investigated in all plant species that were used for comparison of KIPEs3 against MetaCyc, Plant Reactome, and Plant Metabolic Network. As basis for the gene expression served the *A. thaliana* TAIR10 sequence combined with the Araport11 annotation [50], *Cicer arietinum* v1.0 (phytozome 492) [51], *Citrus sinensis* v1.1 (phytozome 154) [52], *Solanum lycopersicum* (phytozome 514 ITAG3.2) [53], and *Daucus carota* v2.0 (phytozome 388) [43]. All available RNA-seq data sets were retrieved from the Sequence Read Archive (SRA) via fastq-dump [54] (Additional File 1).

## Results and Discussion

### Extended bait sequence collection

The initial KIPEs version was exclusively focused on annotating the biosynthesis of flavonoid aglycones [12]. Now, KIPEs3 is able to identify additional candidates that are relevant for the decoration of flavonoid aglycones. The basis for the identification of candidates in new data sets are bait sequences that are previously characterized sequences from other species. The initial KIPEs bait sequence collection was extended by adding anthocyanidin 3-O glucosyltransferase (UGT79B2 and UGT79B3), anthocyanin 5-O-glucosyltransferase (UGT75C1), flavonoid 3-O-glucosyltransferase (UGT78D1 and UGT78D2), UGT72L1, UGT80B1/TT15, anthocyanin acyl transferase (AAT), anthocyanin O-methyl transferase (OMT), BAHD acyltransferase, glutathione S-transferase (GST)-like ligandins/TT19, TT10, anthocyanin transporting ABC transporter, anthocyanin and PA precursor transporting MATE, P _3A_^-^ ATPase AHA10/TT13, and GFS9/TT9 (**Table 1**).

**Table 1:** New flavonoid biosynthesis steps in the KIPEs3 repertoire. These steps were added during the development of KIPEs3. Steps of the core flavonoid aglycone biosynthesis were part of the initial KIPEs release [12]. Required bait sequences and related information are available via GitHub [55].

| Name of the step | Label in KIPes3 | Representative sequence | Reference |
| --- | --- | --- | --- |
| anthocyanidin 3-O glucosyltransferase (UGT79B2 and UGT79B3) | A3GT | AT4G27560 | [24] |
| anthocyanin 5-O-glucosyltransferase (UGT75C1) | A5GT | AT4G14090 | [13] |
| flavonoid 3-O-glycosyltransferase (UGT78D2 and UGT78D1) | F3GT | AT5G17050 | [13] |
| UGT72L1 | UGT72L1 | XP_013443944.2 | [25] |
| UGT80B1/TT15 | TT15 | AT1G43620 | [26] |
| anthocyanidin 3-O-glucoside coumaroyl CoA transferase | 3AT | AT1G03940 | [22] |
| anthocyanidin 5-glucoside malonyl CoA acyl transferase | 5MAT | AT3G29590 | [22] |
| anthocyanidin 3-glucoside malonyl CoA acyl transferase | 3MAT | AAO12206.1 | [27] |
| anthocyanin O-methyl transferase | OMT | VIT_01s0010g03510 | [28] |
| glutathione S-transferase (GST)-like ligandins/TT19 | TT19 | AT5G17220 | [29, 30] |
| Flavonoid oxidase LAC15/TT10 | TT10 | AT5G48100 | [31] |
| anthocyanin transporting ABC transporter | ABCC | AT2G34660 | [32] |
| anthocyanin and PA precursor transporting MATE | MATE | AT3G59030 | [33] |
| $P_{3A^-}$ ATPase AHA10/TT13 | AHA10 | AT1G17260 | [34, 35] |
| GFS9/TT9 | TT9 | AT3G28430 | [36] |

### Major technical updates

The initial Python2 implementation of KIPEs [12] was converted into a Python3 implementation. Syntax changes and the replacement of modules were an integral part of this transformation process. This technical upgrade ensures that more users are able to run KIPEs without installing an outdated Python version. It is also important for long term preservation of KIPEs and to avoid security issues. While upgrading the code to the latest Python version, we also added new features.

The established KIPEs components form the basis of KIPEs3 starting with the identification of orthologs for all bait sequences in the new species of interest (**Figure 1**). Each step in the pathway (enzyme/gene) is represented by one FASTA file containing numerous bait sequences from different plant species. Each collection of bait sequences is used to identify orthologs, i.e. most likely candidates, in the species of interest. This approach is based on the established assumption that orthologs are likely to have the same function (see [12] for details). This identification of orthologs is now based on the widely used Python module dendropy [56]. However, KIPEs3 is not restricted to finding orthologs, but checks a larger set of homologs for their overall sequence similarity and the presence of functionally relevant amino acids. The function of an enzyme is not determined by overall sequence similarity, but can depend on a few essential amino acid residues.

**Figure 1:**
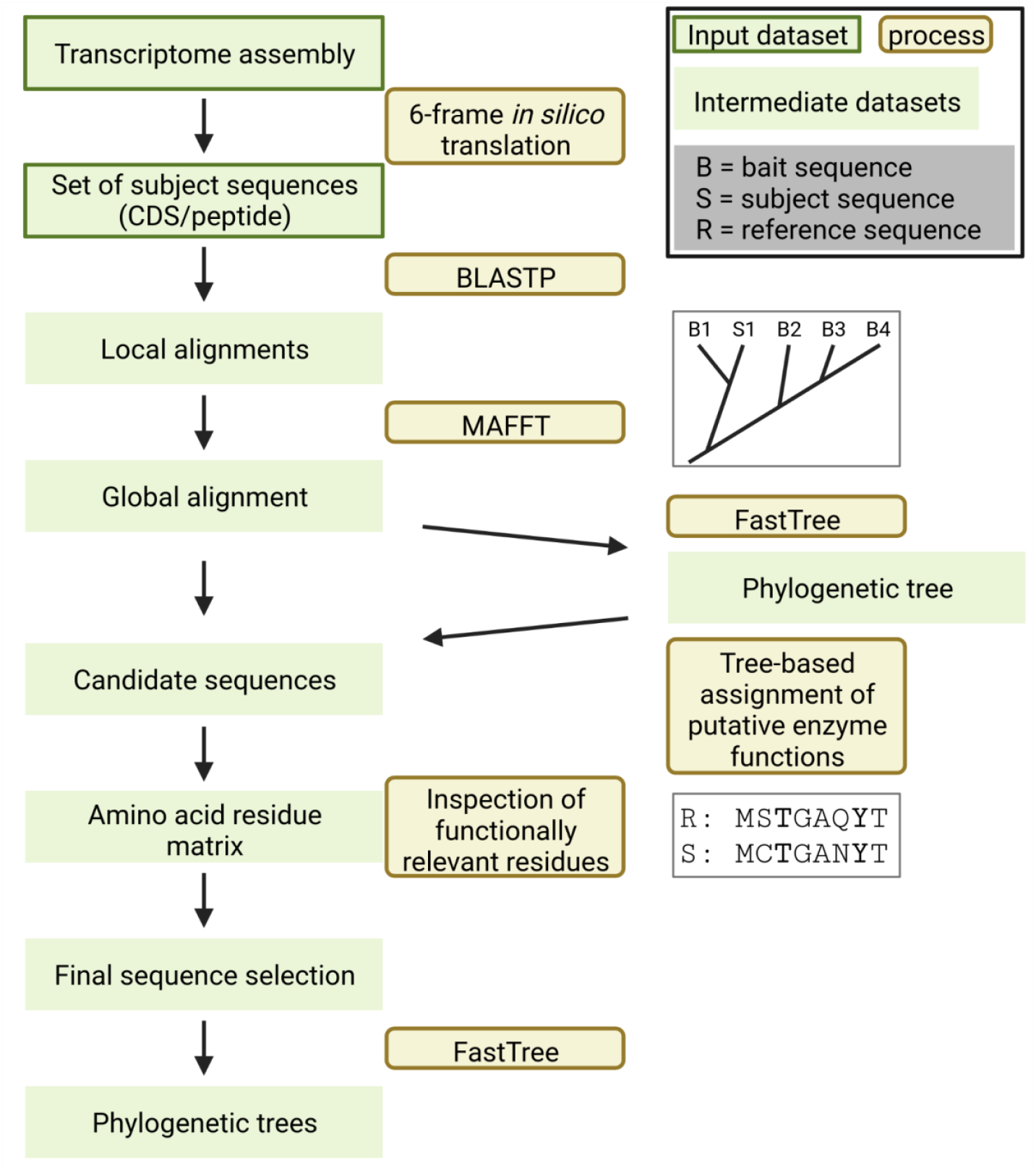
Simplified illustration of central KIPEs3 steps and functions that are included in the standalone and in the online version. Transcriptome or peptide sequences are supplied by the user. Initial candidates are identified by BLAST and checked through global alignments, phylogenetic trees, and the inspection of functionally relevant amino acid residues. Phylogenetic trees are constructed for all final candidates.

A new option allows users to provide expression data that are subjected to an all-vs-all co-expression analysis among the identified candidates. It is generally expected that genes associated with following or preceding steps of the same branch in a biosynthesis pathway show a similar expression pattern, i.e. co-expression with other genes in this pathway branch. A strong co-expression value can further boost the reliability of identified candidates. KIPEs3 will produce a file that can serve as Cytoscape [57] input to visualize the gene expression network (**Figure 2**). This co-expression feature can also help to identify the most important candidate if multiple close paralogs exist for a given function.

**Figure 2:**
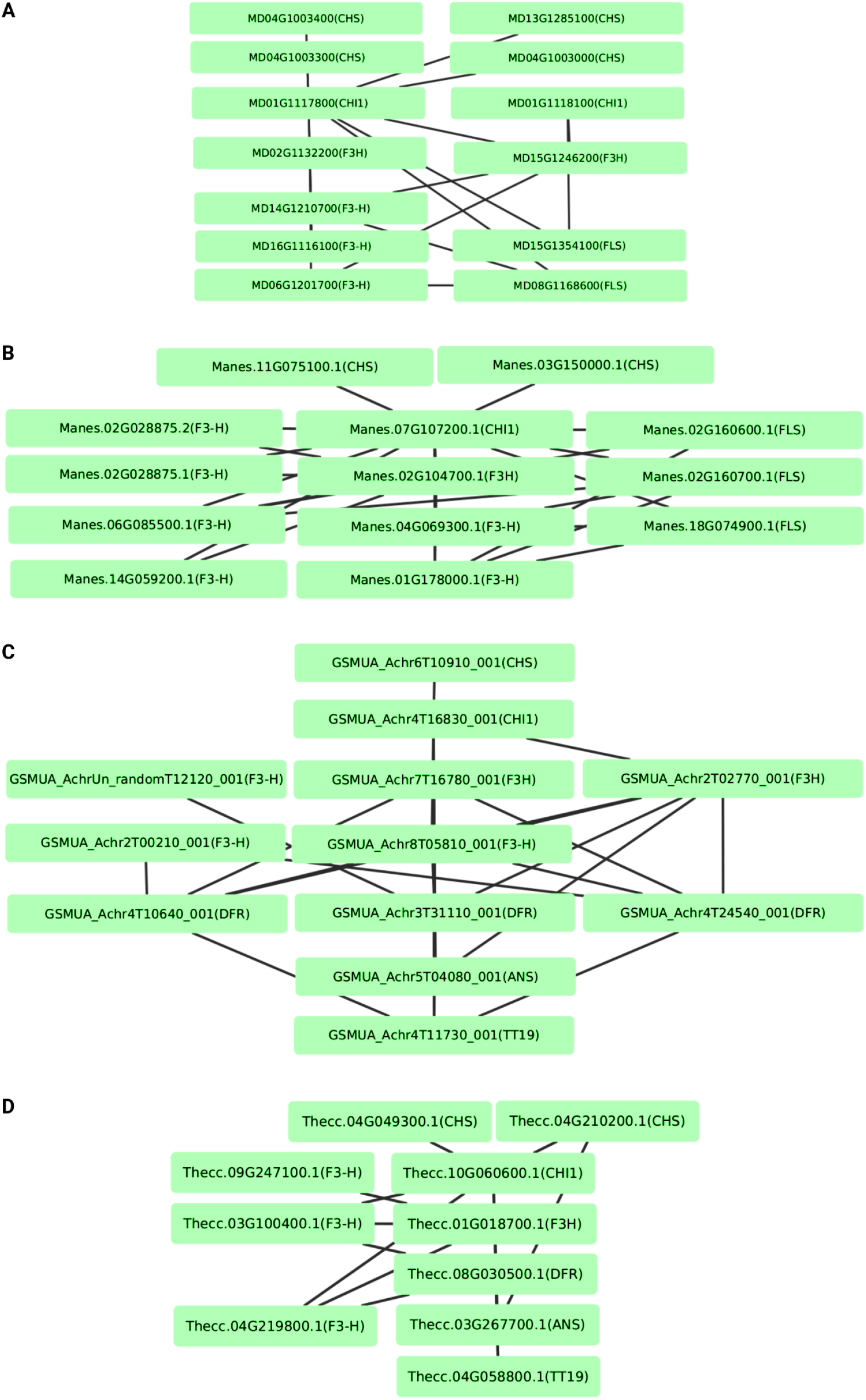
Visualization of co-expression networks via Cytoscape. Flavonol biosynthesis genes in *Malus domestica* (A) and *Manihot esculenta* (B) were analyzed based on co-expression. The anthocyanin biosynthesis genes were studied in *Musa acuminata* (C) and *Theobroma cacao* (D).

Many KIPEs users expressed an interest in phylogenetic trees that place the identified candidates in the context of previously characterized homologous sequences. These trees can help to explore the relationships of multiple candidates to previously characterized bait or reference sequences. KIPEs3 can automatically construct trees of the identified candidates and the corresponding bait sequences for all steps in the pathway. This enables a manual inspection without the need for additional phylogenetic analyses. KIPEs3 generates files in the Newick format which can be visualized and modified for publication with established tools like FigTree or iTOL [58].

KIPEs is able to start with coding sequences or with peptide sequences. It is even possible to provide a genome sequence, but performing the gene prediction with dedicated tools is recommended for most applications. Generally, it is best to provide peptide sequences as this speeds up the analysis. Users define the type of input sequences that they are providing. Since this information might not always be accurate, an option was added to check if the provided sequences comprise amino acids or nucleotides. A warning is shown if the result of this analysis contradicts the sequence type set by the user. However, it remains unfeasible to distinguish a collection of coding sequences from a highly fragmented genome sequence assembly based on short reads. Therefore, it remains the users’ responsibility to set the correct sequence type.

Given the power of modern hardware, the computational resources required for running KIPEs are neglectable. A single core and less than 2GB of memory are already sufficient for the analysis. The entire analysis is completed within minutes, but the precise run time depends on the specific settings and size of the analyzed data set (**Figure 3**). Especially large data sets can cause run times of about one hour, but most plant species can be analyzed in less than 20 minutes (**Figure 3A**). This run time analysis was performed on Phytozome data sets of a wide range of plant species to simulate a realistic application of KIPEs3.

**Figure 3:**
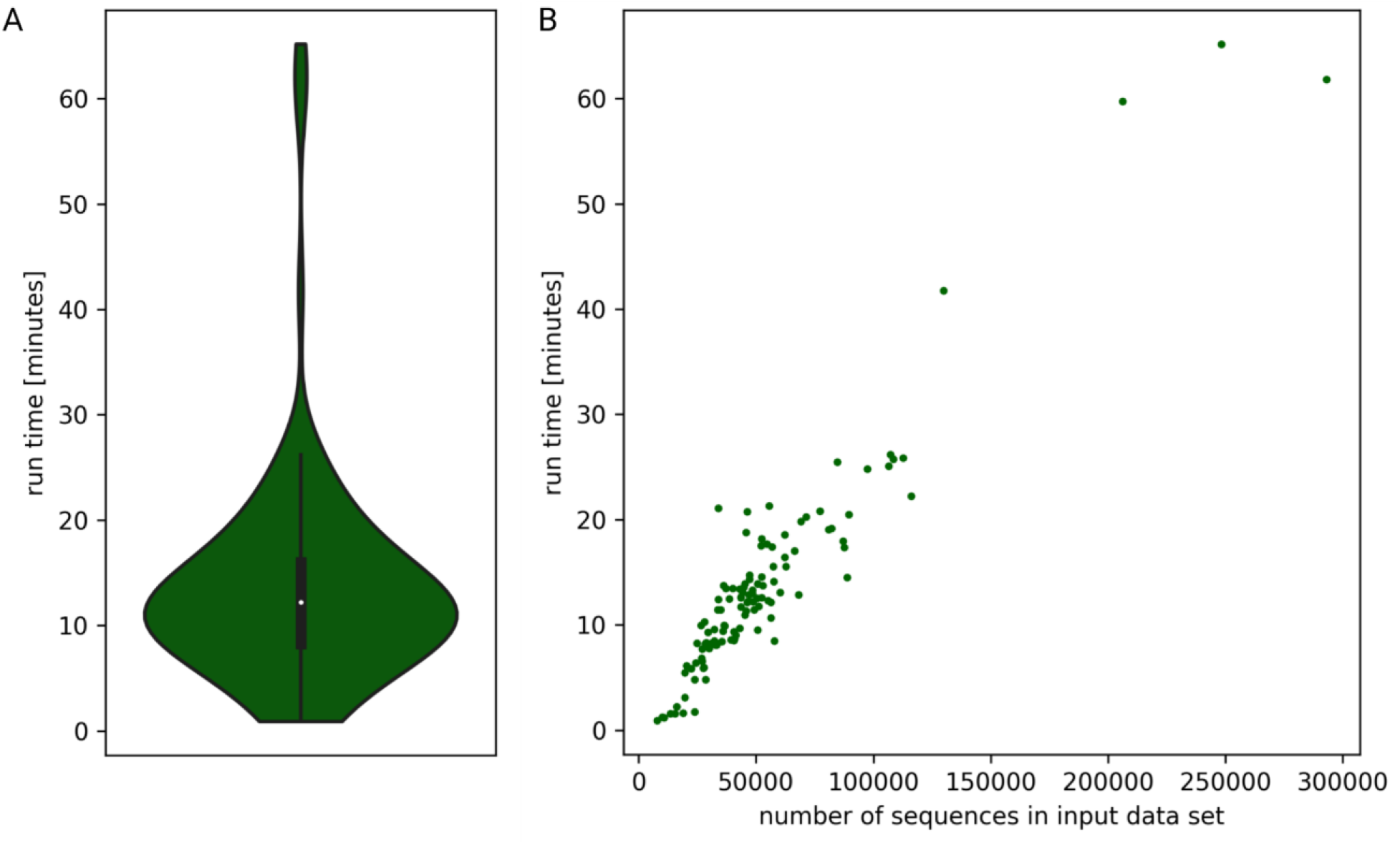
KIPEs3 run time analysis based on 121 Phytozome data sets (A) revealed that the run time is depending on the number of sequences in the input data set (B).

### Comparison of KIPEs3 to other databases

MetaCyc, Plant Metabolic Network (PMN), and Plant Reactome are comprehensive databases describing the pathways, enzymes, and substrate compounds. While MetaCyc contains information about all domains of life, PMN and Plant Reactome are specific to plants and green algae. KIPEs3 is a specialized tool which provides individual gene level annotation for a user defined biosynthesis pathway in any species. MetaCyc contains only genes with experimentally determined functions in the represented pathways. PMN comprises 126 species-specific datasets and houses PlantCyc which covers over 500 additional plant species. PlantCyc harbors experimentally supported and predicted gene functions. The predicted enzymes are classified using the Ensemble Enzyme Prediction Pipeline (E2P2) software and RPSD v4.2 [59]. E2P2 uses BLAST [60] and PRIAM [61] to assign functions using sequence similarity to previously-known genes sourced from MetaCyc, BRENDA [62], and SwissProt [63] among others. The genome sequences used for this annotation are selected on the basis of at least 75% BUSCO (Benchmarking Universal Single-Copy Orthologs) completeness [34]. Plant Reactome uses *Oryza sativa* as a reference species to predict enzymes in 120 other species currently available in the database. Enzyme prediction in a species is based on orthologs of that enzyme in rice. Enzyme prediction in KIPEs3 is more stringent as it starts with ortholog identification based on multiple experimentally identified genes defined in literature (bait sequences). Ortholog identification is followed by an in-depth sequence similarity analysis including a check of the presence of functionally relevant amino acids. Only those candidates which pass all these steps are reported as most likely functional. Moreover, KIPEs3 can also identify similar candidates that do not fulfill all selection criteria. Contrary to most databases, researchers can use their own assembled dataset when running KIPEs3. This option enables the functional annotation of new genome sequences to identify a particular pathway of interest, given some prior knowledge about the pathway. Some features of these databases compared to KIPEs are summarized in **Table 2**.

**Table 2:**
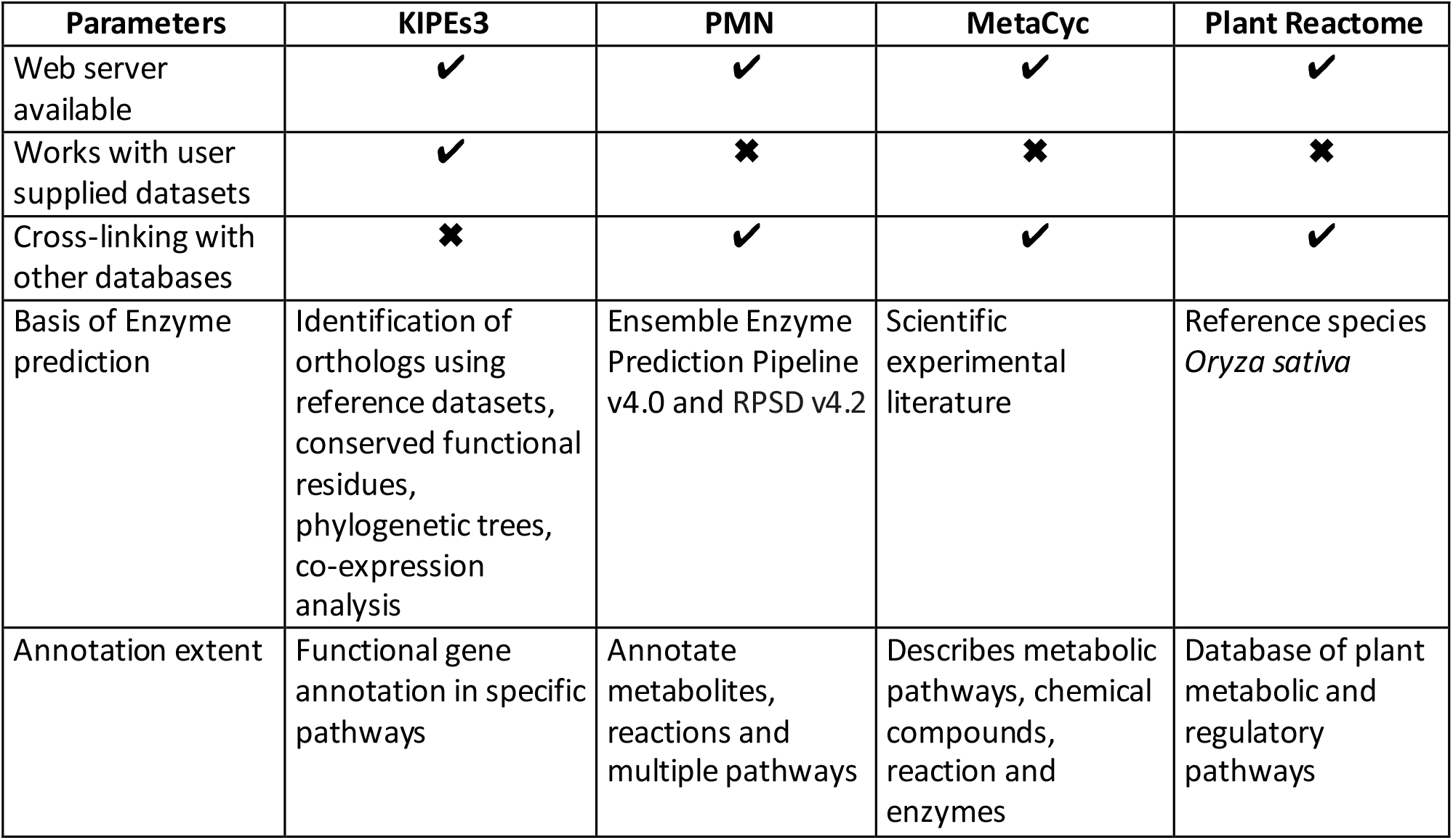
Comparison of KIPEs3 against established pathway databases.

### Functional annotation performance of KIPEs3 compared to other databases

We compared the annotation results for FLS and LYC-b in terms of correctness of the annotation. Functional annotation correctness of a candidate can be assessed (a) in terms of correct amino acid residues required for a functional enzyme, (b) in terms of its position in a phylogenetic tree with characterized orthologous sequences, and (c) in terms of expression pattern. A higher gene expression is usually suggesting functional relevance of a given candidate. Co-expression with other members of the same biosynthesis pathways is also helpful in identifying the most promising candidates. We followed a combined approach for LYC-b and FLS annotation. The annotation correctness was assessed by the co-expression of a given candidate with other genes of the flavonoid or carotenoid biosynthesis, respectively. The presence of functionally relevant amino acid residues served as the final and deciding factor for declaring a candidate gene as “functional”.

Lycopene β cyclase (LYC-b) is one of the key enzymes in the carotenoid biosynthesis pathway and catalyzes the β-cyclization of both ends of lycopene to produce β-lycopene. LYC-b is known to have conserved domains such as “dinucleotide binding motif”, “Cyclase motif I and II”, “Conserved region β-LYC”, and a “β cyclase motif” [64]. Since studies about the amino acid residues responsible for catalytic sites are still lacking, a phylogenetic tree of LYC-b candidates was constructed to identify likely functional orthologs. The carotenoid genes, neoxanthin synthase (NSY) and lycopene ε-cyclase (LYC-ε), were taken as an outgroup due to their high structure similarity to LYC-b [65] (Additional File 2, Additional File 3).

FLS is a key enzyme of the flavonol biosynthesis pathway and catalyzes the oxidation of dihydroflavonols to flavonols. FLS belongs to the 2-oxoglutarate dependent dioxygenase (2-ODD) superfamily. They are characterized by conserved 2-oxoglutarate binding residues (Arg287 and Ser289) and conserved ferrous iron-binding residues (His221, Asp223, and His277) [66]. A functional FLS also possesses residues responsible for proper folding of 2-ODD polypeptide (Gly68, His75, Pro207, and Gly261) [67]. FLS specific motifs, “PxxxIRxxxEQP” and “SxxTxLVP”, distinguish FLS from other plant 2 -ODDs like F3H, ANS, and FNS II (Additional File 4).

We used two metrics (*specificity* and *accuracy*) to characterize the annotation performance of the different tools and databases. *Specificity* describes how well the database/tool can annotate a functional enzyme. It is calculated as the ratio of the number of correct candidates predicted by the database/tool to the total number of functional candidates present in that species. *Accuracy* denotes the ability of a database/tool to discriminate between functional and non-functional candidates. It is calculated as the ratio of correct predictions to the total predictions made by a database/tool for a given species and a gene in a pathway. Finally, the annotation performance is calculated as the average sum of s*pecificity* and *accuracy*. For LYC-b annotation, we found that the performance of Plant Reactome and KIPEs was 100% in all three species, *Arabidopsis*, tomato, and carrot. MetaCyc showed 100% annotation performance for *Arabidopsis*, 75% for tomato, and dropped to 0% for carrot as it did not annotate any LYC-b gene in carrot. Despite the high specificity (100%) of Plant Metabolic Network, the performance was between 65-75% for all three species due to low accuracy. (**Figure 4 (a-f)**). KIPEs3 maintained its 100% performance in all three species, *Arabidopsis*, chickpea, and orange, used for FLS annotation as well. Like KIPEs3, MetaCyc provides a correct *Arabidopsis* FLS annotation. However, MetaCyc does not report any FLS in chickpea and orange, while KIPEs3 identified candidates in both species. Plant Reactome was neither specific nor accurate in FLS annotation and failed to annotate any candidate in any of the above species. Plant Metabolic Network showed low performance (∼0-60%) again due to low accuracy (**Figure 4 (g-i)**).

**Figure 4:**
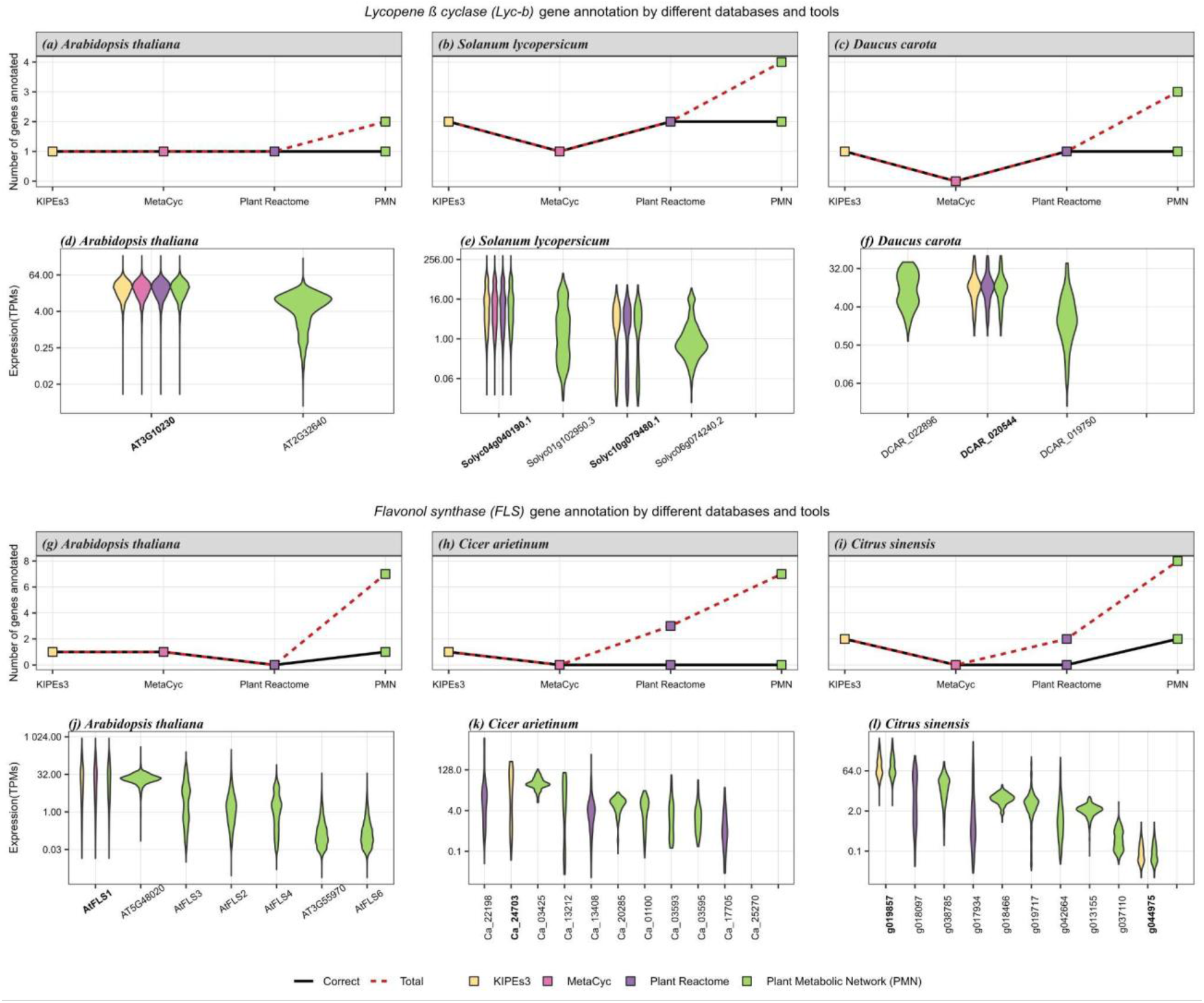
Functional annotation tool comparison using the *LYC-b* gene from the carotenoid biosynthesis pathway and the *FLS* gene from the flavonoid biosynthesis pathway. Comparison of LYC-b annotation **(a-f)** between KIPEs and three databases. The number of annotated (red dashed line) and correctly annotated (black solid line) LYC-b genes is displayed per tool/database for *Arabidopsis thaliana* **(a)**, *Solanum lycopersicum* **(b)**, and *Daucus carota* **(c)**. The corresponding gene expression (TPM) of all candidate genes was investigated to differentiate between genes and potential pseudogenes **(d-f)** for the three species, respectively. Functional gene IDs are written in *boldface*. Comparison of FLS annotation **(g-l)** between KIPEs and three databases. The number of annotated (red dashed line) and correctly annotated (black solid line) FLS genes is displayed per tool/database for *Arabidopsis thaliana* **(g)**, *Cicer arietinum* **(h)**, and *Citrus sinensis* **(i)**. The corresponding gene expression (TPM) of all candidate genes was investigated to differentiate between genes and potential pseudogenes **(j-l)** for the three species, respectively. Functional gene IDs are written in *boldface*.

Plant Metabolic Network exhibited the least accuracy which is likely due to the function prediction approach using sequence similarity alone which can lead to inaccurate prediction. Highly similar enzymes belonging to the same family might have similar sequences, but may have different functions *in vivo*. The performance of Plant Reactome was also low in case of FLS annotation emphasizing the downfall of using only rice in curation and prediction, as genes in species distant from rice will be predicted less reliably and genes missing in rice cannot be predicted in other species. MetaCyc contains only experimentally evidenced genes, hence has high accuracy but low specificity. This also leads to annotation of a limited number of genes and species.

In summary, KIPEs3 showed the best performance for functional annotation of genes having both high accuracy as well as specificity. This makes KIPEs3 a good tool for annotating novel genome sequences with focus on a particular pathway with a substantial amount of prior knowledge about the involved genes in other species.

### KIPEs3 is available as an online service

KIPEs3 is provided as an online service hosted on a webserver at TU Braunschweig [68]. Users can upload their FASTA files containing coding sequences or peptide sequences of a species. Bait sequence sets are stored on the server and can be directly used without the need to be manually uploaded by the user. Optional parameters are pre-configured to allow an automatic analysis with default settings, but can also be modified by the user through the web interface if necessary. Users may opt-in to receive email notifications once their analysis is finished and the results are available. Alternatively, users may choose to store a cookie to allow their browser to keep track of the progress. The archive provided for download includes all result files that users would get when running KIPEs3 locally.

The web server is implemented in Python3 using the Django web framework v4 [69]. Whenever a user uploads a file and submits a new job through the form on the website, the form data is sent to the server via an HTTP POST request and the file is temporarily stored on the server machine. The job is placed into a queue and executed as soon as there are enough free resources (disk space, memory, and CPU cores) available. KIPEs3 and all its dependencies are installed on the server machine. The user is redirected to a page with a console showing the command that is executed, and the program output is displayed in real time using XMLHttpRequests. All user supplied data are deleted from the server 48 hours after the job has been finished and the results have been downloaded.

### Analysis of the flavonoid biosynthesis in *A. thaliana* Nd-1, *Capsella rubella*, and *Dioscorea dumetorum*

First, a technical check was performed by analyzing the flavonoid biosynthesis in the *A. thaliana* accession Niederzenz-1 (Nd-1). It is expected that orthologs of all characterized genes of the reference accession Columbia-0 (Col-0) are also present in Nd-1, while the sequences are slightly different from the bait sequences. All genes known in Col-0 were also discovered in Nd-1 (Additional File 2). Next, *Capsella grandiflora* was analyzed with KIPEs3 to show that the identification of flavonoid biosynthesis genes works also in species that are not represented in most of the bait sequence sets like *A. thaliana*. Genes present in *A. thaliana* were also discovered in the close relative *C. grandiflora* (Additional File 3). To demonstrate the potential of KIPEs3 to annotate the flavonoid biosynthesis in a distant relative of *A. thaliana*, the flavonoid biosynthesis of *Dioscorea dumetorum*, which belongs to the monocots, was analyzed (Additional File 5).

The new flavonoid biosynthesis pathway steps added to the repertoire of KIPEs3 include the diverse and promiscuous decoration enzymes like glycosyltransferases, acyltransferases, and methyltransferases. Since some of these steps might be species-specific, it is possible that some steps are missing in a given plant species of interest. This requires specific attention when interpreting the KIPEs3 results to avoid false positives. While the aglycon biosynthesis is highly conserved and catalyzed by well characterized enzymes, flavonoid decorating enzymes are less studied and show a high level of promiscuity [70–72]. This poses a challenge for the identification through sequence similarity. Little is known about the functionally relevant amino acids in the flavonoid decorating enzymes. In addition, this extension of the KIPEs3 flavonoid biosynthesis bait sequence sets is no longer restricted to enzymes, but also includes flavonoid transporters. They do not have an active center that could be composed of functionally relevant amino acids. This turns the reliable identification of transport-associated proteins into a remaining challenge.

### Analysis of the carotenoid biosynthesis

A novel set of bait sequences was compiled to enable the annotation of the carotenoid biosynthesis with KIPEs3 (**Table 3**). The carotenoid biosynthesis is well studied in many plant species including *A. thaliana* [73]. The set of compiled bait and reference sequences was evaluated by running the annotation workflow on *A. thaliana* Nd-1, *Daucus carota, C. rubella*, and *D. dumetorum* (**Figure 5**; Additional File 3). *Daucus carota* carotenoid biosynthesis genes previously described in the literature [74] were recovered by KIPEs3 (Additional File 6). This supports the accuracy of the candidate gene identification.

**Table 3:** Carotenoid biosynthesis genes. ^1^ most likely non-functional ^2^ BCH duplication after monocot-dicot split

| EC Number | Name of the step | Representative sequence / Label in KIPes3 | Reference |
| --- | --- | --- | --- |
| EC:2.5.1.32 | Phytoene synthase (PSY) | AT5G17230 | [75] |
| EC:1.3.5.5 | Phytoene desaturase (PDS) | AT4G14210 | [76] |
| EC:1.3.5.6 | Zeta-carotene desaturase (ZDS) | AT3G04870 | [77] |
| EC:5.2.1.12 | 15-cis-zeta-carotene isomerase (ZISO) | AT1G10830 | [78] |
| EC:5.2.1.13 | Carotenoid isomerase 1 (CRTISO1/CCR2) | AT1G06820 | [79] |
| ----- | Carotenoid isomerase 2 (CRTISO2)<sup>1</sup> | AT1G57770 | - |
| EC:5.5.1.19 | Lycopene cyclase (LCYB/CRTL-B) | AT3G10230 | [80] |
| EC:5.5.1.18 | Lycopene cyclase (LCYE/CRTL-E) | AT5G57030 | [81] |
| EC:1.14.15.24 | Beta-carotene hydroxylase 1 (BCH1/CHYB1/CHY1) | AT4G25700 | [82] |
| EC:1.14.15.24 | Beta-carotene hydroxylase 2 | AT5G52570 | [83] |
|  | (BCH2/CHYB2/CHY2)<sup>2</sup> |  |  |
| EC:1.14.-.- | Carotene beta-ring hydroxylase (CYP97A3/LUT5) | AT1G31800 | [84] |
| EC:1.14.-.- | Cytochrome P450 97B3 (CYP97B3) | AT4G15110 | [85] |
| EC 1.14.14.158 | Carotene epsilon-monooxygenase (CYP97C1/CHYE/LUT1) | AT3G53130 | [86] |
| EC 1.14.15.21 | Zeaxanthin epoxidase (ZEP/ABA1/NPQ2) | AT5G67030 | [87] |
| EC 1.23.5.1 | Violaxanthin de-epoxidase (VDE/NPQ1) | AT1G08550 | [87] |
| EC 5.3.99.9 | Neoxanthin synthase (NSY/ABA4) | AT1G67080 | [88] |

**Figure 5:**
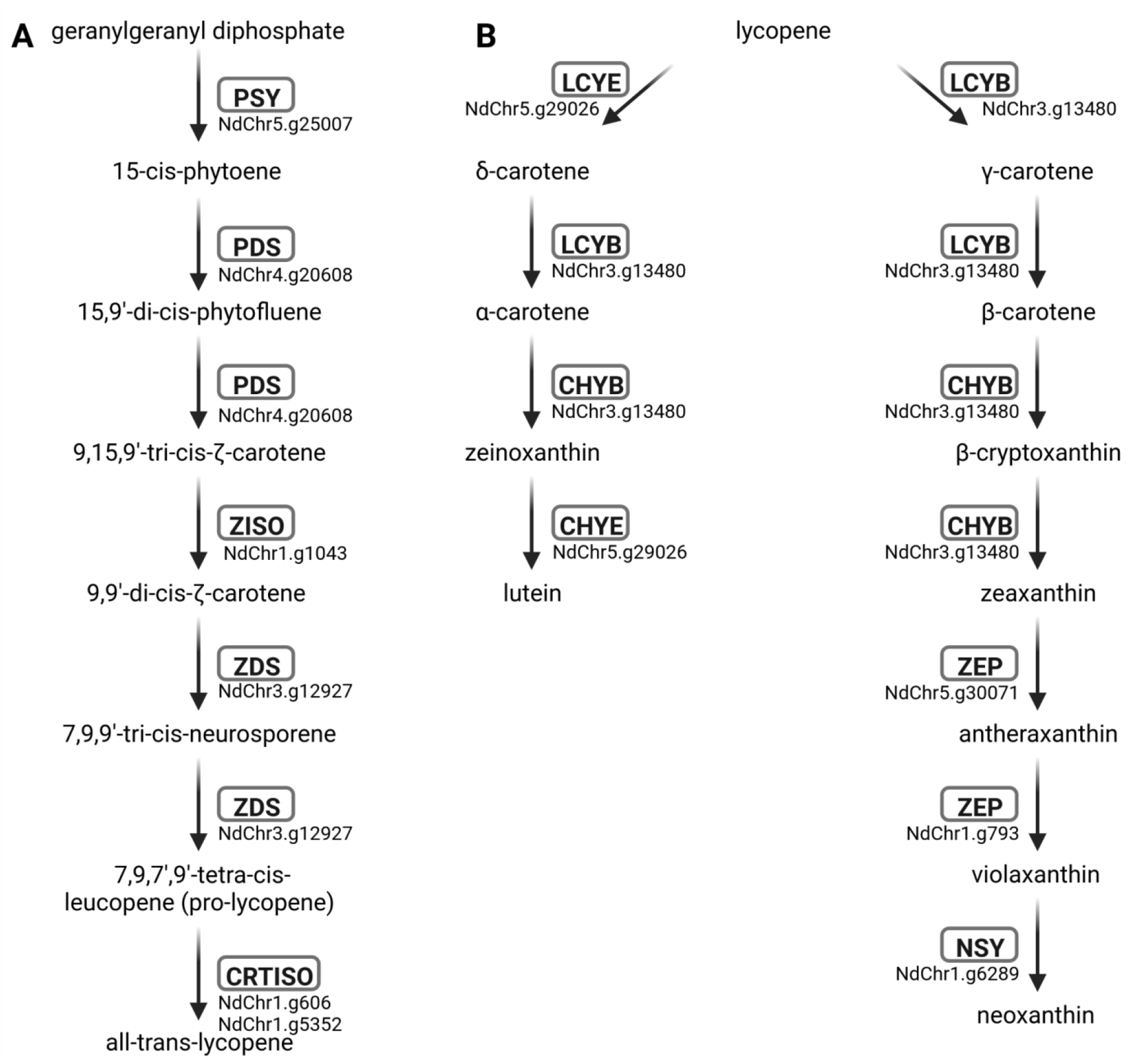
Carotenoid biosynthesis genes in *Arabidopsis thaliana* Nd-1. The carotenoid biosynthesis comprises a linear part (A) and a branching part (B).

### Transcriptional regulation of biosynthesis pathways

Many plant biosynthesis pathways are regulated at the transcriptional level by members of large transcription factor families. Examples are the R2R3-MYBs and bHLHs that form complexes involved in the control of the flavonoid biosynthesis [89]. As we demonstrated previously, it is possible to analyze large gene families like MYBs and bHLHs with KIPEs [12]. However, KIPEs3 is optimized for the analysis of pathways that comprise genes with low copy numbers. Most structural genes in these pathways are part of a small gene family. Large gene families like the P450 cytochromes (CYPs) or 2-oxoglutarate dioxygenases (2-ODDs) comprise multiple smaller subfamilies with distinct biological functions. Only the individual subfamilies are relevant for the analysis with KIPEs3. Ideally, the application of KIPEs3 should be restricted to the annotation of structural genes in a biosynthesis. While it is possible to investigate large transcription factor families, dedicated tools will perform better. Members of the MYB and bHLH gene families can be annotated by specific tools [47]. General phylogeny-based approaches can be deployed to study other involved transcription factor families [90, 91].

## Conclusions

While the generation of high-quality genome sequences is turning into a routine task, the structural and especially the functional annotation of these sequences is an increasing challenge. The ability to annotate biosynthesis pathways beyond the flavonoid biosynthesis positions KIPEs3 as a versatile platform for functional annotation in projects focused on specialized metabolites. We demonstrated the capabilities of KIPEs3 by comparing the annotation of *FLS* and *LYC-b* in three species against the results of other popular tools and databases. We showed that the use of broad taxonomic range bait sequences and functionally relevant amino acids allows KIPEs3 to recover more correct annotations per pathway than other tools that rely on general sequence similarity. KIPEs3 can easily annotate novel genome sequences with focus on a particular pathway while this is not possible with other annotation tools. Although we focused on plant metabolism in this study, KIPEs3 could also be applied for the investigation of non-plant organisms like fungi or bacteria. A sufficient amount of prior knowledge about the pathway in other species is the only requirement. The automatic annotation of biosynthesis pathways paves the way for comparative genomic studies that are looking for species-specific differences or intraspecific variation.

## Availability and requirements

Project name: KIPEs3

Project home page: https://github.com/bpucker/KIPEs

Operating system(s): Linux (website is platform independent)

Programming language: Python3

Other requirements: BLAST, MAFFT, FastTree2, dendropy, scipy

License: GNU General Public License v3.0

RRID: SCR_022370

## Data Availability

All data sets analyzed in this study are publicly available. Data sets generated as part of this study are shared via GitHub (https://github.com/bpucker/KIPEs). A docker image is available via DockerHub (https://hub.docker.com/r/bpucker/kipes).

## Declarations

### List of abbreviations

Not applicable

### Ethics approval and consent to participate

Not applicable

### Consent for publication

Not applicable

## Competing interests

AR and NC have no competing interests. BP is head of the technology transfer center Plant Genomics and Applied Bioinformatics at iTUBS.

## Funding

We acknowledge support by the Open Access Publication Funds of Technische Universität Braunschweig.

## Authors’ contributions

AR developed the web server framework and contributed to the manuscript. NC performed benchmarking of KIPEs3 and improved the carotenoid biosynthesis annotation. BP designed the research, conducted experiments, wrote the first draft, and revised the manuscript.

## Acknowledgements

Many thanks to the German network for bioinformatics infrastructure (de.NBI, grant 031A533A) and the Bioinformatics Resource Facility (BRF) at the Center for Biotechnology (CeBiTec) at Bielefeld University for providing an environment to perform the computational analyses. bioRender.com was used to construct some of the figures.

## Additional files

**Additional file 1**

Accession numbers of the analyzed RNA-seq datasets.

**Additional file 2**

Multiple sequence alignment of the LYC-b candidates in three species (*Arabidopsis thaliana, Solanum lycopersicum*, and *Daucus carota*) annotated by KIPEs3, MetaCyc, Plant Reactome, and Plant Metabolic Network. The alignment was generated using MAFFTv7. Characteristic regions of plant LCY-bs are indicated above the sequences: Dinucleotide binding site, Cyclase motifs (CM) I and II, and Charged region. Domains described as essential for LCY-b activity are shown as LCY-b CAD (Catalytic Activity Domain). The functional LYC-b on the basis of presence of conserved amino acid residues are highlighted in blue color.

**Additional file 3**

Phylogenetic tree of the LYC-b candidates in the three species (*Arabidopsis thaliana, Solanum lycopersicum*, and *Dacus carota*). The highly similar sequences of Neoxanthin synthase (NSY) and Lycopene-e-cylase (LYC-e) were included as outgroup (highlighted in gray). The functional LYC-b clade was highlighted in the phylogenetic tree.

**Additional file 4**

Multiple sequence alignment of the FLS in three species (*Arabidopsis thaliana, Cicer arietinum*, and *Citrus sinensis*) annotated by KIPEs3, MetaCyc, Plant Reactome, and Plant Metabolic Network. The alignment was generated using MAFFTv7. The two black boxes highlight the FLS-specific motifs “PxxxIRxxxEQP” and “SxxTxLVP.” Amino acid residues responsible for binding ferrous iron (H221, D223, and H277) and 2-oxoglutarate (R287 and S289) are marked with black and gray arrows, respectively. The residues known to be involved in the proper folding of the 2-ODD polypeptide are marked with asterisks (G68, H75, P207, and G261). The functional FLS on the basis of the presence of conserved amino acid residues are highlighted in red-yellow color.

**Additional file 5**

Results of the flavonoid biosynthesis analysis in *Arabidopsis thaliana* Nd-1, *Capsella rubella*, and *Dioscorea dumetorum*.

**Additional file 6**

Results of the carotenoid biosynthesis analysis in *A. thaliana* Nd-1, *Daucus carota, C. rubella*, and *D. dumetorum*.

## Notes

https://github.com/bpucker/KIPEs

https://hub.docker.com/r/bpucker/kipes

https://pbb-tools.de/

### Summary of Updates

web server URL updated

## References

1. Young AJ. The photoprotective role of carotenoids in higher plants. Physiol Plant. 1991;83:702–8.

2. Frank HA, Cogdell RJ. Carotenoids in Photosynthesis. Photochem Photobiol. 1996;63:257–64.

3. Polívka T, Frank HA. Molecular factors controlling photosynthetic light harvesting by carotenoids. Acc Chem Res. 2010;43:1125–34.

4. Moehs CP, Tian L, Osteryoung KW, DellaPenna D. Analysis of carotenoid biosynthetic gene expression during marigold petal development. Plant Mol Biol. 2001;45:281–93.

5. Wan H, Yu C, Han Y, Guo X, Luo L, Pan H, et al. Determination of Flavonoids and Carotenoids and Their Contributions to Various Colors of Rose Cultivars (Rosa spp.). Front Plant Sci. 2019;10.

6. Wang Y, Zhang C, Dong B, Fu J, Hu S, Zhao H. Carotenoid Accumulation and Its Contribution to Flower Coloration of Osmanthus fragrans. Front Plant Sci. 2018;9.

7. Parry AD, Horgan R. Carotenoids and abscisic acid (ABA) biosynthesis in higher plants. Physiol Plant. 1991;82:320–6.

8. Dudek B, Warskulat A-C, Schneider B. The Occurrence of Flavonoids and Related Compounds in Flower Sections of Papaver nudicaule. Plants. 2016;5:28.

9. Schiestl FP, Johnson SD. Pollinator-mediated evolution of floral signals. Trends Ecol Evol. 2013;28:307–15.

10. Stavenga DG, Leertouwer HL, Dudek B, van der Kooi CJ. Coloration of Flowers by Flavonoids and Consequences of pH Dependent Absorption. Front Plant Sci. 2021;11.

11. Winkel-Shirley B. Flavonoid Biosynthesis. A Colorful Model for Genetics, Biochemistry, Cell Biology, and Biotechnology. Plant Physiol. 2001;126:485–93.

12. Pucker B, Reiher F, Schilbert HM. Automatic Identification of Players in the Flavonoid Biosynthesis with Application on the Biomedicinal Plant Croton tiglium. Plants. 2020;9:1103.

13. Tohge T, Nishiyama Y, Hirai MY, Yano M, Nakajima J, Awazuhara M, et al. Functional genomics by integrated analysis of metabolome and transcriptome of Arabidopsis plants over-expressing an MYB transcription factor. Plant J Cell Mol Biol. 2005;42:218–35.

14. Slámová K, Kapešová J, Valentová K. “Sweet Flavonoids”: Glycosidase-Catalyzed Modifications. Int J Mol Sci. 2018;19:2126.

15. Khodzhaieva RS, Gladkov ES, Kyrychenko A, Roshal AD. Progress and Achievements in Glycosylation of Flavonoids. Front Chem. 2021;9.

16. Grotewold E, Davies K. Trafficking and Sequestration of Anthocyanins. Nat Prod Commun. 2008;3:1934578X0800300806.

17. Zhao J, Dixon RA. The “ins” and “outs” of flavonoid transport. Trends Plant Sci. 2010;15:72–80.

18. Pucker B, Selmar D. Biochemistry and Molecular Basis of Intracellular Flavonoid Transport in Plants. Plants Basel Switz. 2022;11:963.

19. Mazza G, Brouillard R. Recent developments in the stabilization of anthocyanins in food products. Food Chem. 1987;25:207–25.

20. Tohge T, de Souza LP, Fernie AR. Current understanding of the pathways of flavonoid biosynthesis in model and crop plants. J Exp Bot. 2017;68:4013–28.

21. Li X, Li Y, Zhao M, Hu Y, Meng F, Song X, et al. Molecular and Metabolic Insights into Anthocyanin Biosynthesis for Leaf Color Change in Chokecherry (Padus virginiana). Int J Mol Sci. 2021;22:10697.

22. Luo J, Nishiyama Y, Fuell C, Taguchi G, Elliott K, Hill L, et al. Convergent evolution in the BAHD family of acyl transferases: identification and characterization of anthocyanin acyl transferases from Arabidopsis thaliana. Plant J. 2007;50:678–95.

23. Sakuta M, Tanaka A, Iwase K, Miyasaka M, Ichiki S, Hatai M, et al. Anthocyanin synthesis potential in betalain-producing Caryophyllales plants. J Plant Res. 2021. https://doi.org/10.1007/s10265-021-01341-0.

24. Gerats AGM, Martin C. Flavonoid Synthesis in Petunia Hybrida; Genetics and Molecular Biology of Flower Colour. In: Stafford HA, Ibrahim RK, editors. Phenolic Metabolism in Plants. Boston, MA: Springer US; 1992. p. 165–99.

25. Knoch E, Sugawara S, Mori T, Nakabayashi R, Saito K, Yonekura-Sakakibara K. UGT79B31 is responsible for the final modification step of pollen-specific flavonoid biosynthesis in Petunia hybrida. Planta. 2018;247:779–90.

26. Martin C, Prescott A, Mackay S, Bartlett J, Vrijlandt E. Control of anthocyanin biosynthesis in flowers of Antirrhinum majus. Plant J. 1991;1:37–49.

27. Fujino N, Tenma N, Waki T, Ito K, Komatsuzaki Y, Sugiyama K, et al. Physical interactions among flavonoid enzymes in snapdragon and torenia reveal the diversity in the flavonoid metabolon organization of different plant species. Plant J. 2018;94:372–92.

28. Sparvoli F, Martin C, Scienza A, Gavazzi G, Tonelli C. Cloning and molecular analysis of structural genes involved in flavonoid and stilbene biosynthesis in grape (Vitis vinifera L.). Plant Mol Biol. 1994;24:743–55.

29. Braidot E, Zancani M, Petrussa E, Peresson C, Bertolini A, Patui S, et al. Transport and accumulation of flavonoids in grapevine (Vitis vinifera L.). Plant Signal Behav. 2008;3:626–32.

30. Matus JT, Poupin MJ, Cañón P, Bordeu E, Alcalde JA, Arce-Johnson P. Isolation of WDR and bHLH genes related to flavonoid synthesis in grapevine (Vitis vinifera L.). Plant Mol Biol. 2010;72:607–20.

31. Grotewold E, Drummond BJ, Bowen B, Peterson T. The myb-homologous P gene controls phlobaphene pigmentation in maize floral organs by directly activating a flavonoid biosynthetic gene subset. Cell. 1994;76:543–53.

32. Marks RA, Hotaling S, Frandsen PB, VanBuren R. Representation and participation across 20 years of plant genome sequencing. Nat Plants. 2021;7:1571–8.

33. Pucker B, Irisarri I, Vries J de, Xu B. Plant genome sequence assembly in the era of long reads: Progress, challenges and future directions. Quant Plant Biol. 2022;3:e5.

34. Hawkins C, Ginzburg D, Zhao K, Dwyer W, Xue B, Xu A, et al. Plant Metabolic Network 15: A resource of genome-wide metabolism databases for 126 plants and algae. J Integr Plant Biol. 2021;63:1888–905.

35. Naithani S, Gupta P, Preece J, D’Eustachio P, Elser JL, Garg P, et al. Plant Reactome: a knowledgebase and resource for comparative pathway analysis. Nucleic Acids Res. 2020;48:D1093 –103.

36. Caspi R, Altman T, Billington R, Dreher K, Foerster H, Fulcher CA, et al. The MetaCyc database of metabolic pathways and enzymes and the BioCyc collection of Pathway/Genome Databases. Nucleic Acids Res. 2014;42:D459–71.

37. Jones P, Binns D, Chang H-Y, Fraser M, Li W, McAnulla C, et al. InterProScan 5: genome-scale protein function classification. Bioinformatics. 2014;30:1236–40.

38. Conesa A, Götz S, García-Gómez JM, Terol J, Talón M, Robles M. Blast2GO: a universal tool for annotation, visualization and analysis in functional genomics research. Bioinformatics. 2005;21:3674–6.

39. Moriya Y, Itoh M, Okuda S, Yoshizawa AC, Kanehisa M. KAAS: an automatic genome annotation and pathway reconstruction server. Nucleic Acids Res. 2007;35 Web Server issue:W182–5.

40. Schwacke R, Ponce-Soto GY, Krause K, Bolger AM, Arsova B, Hallab A, et al. MapMan4: A Refined Protein Classification and Annotation Framework Applicable to Multi-Omics Data Analysis. Mol Plant. 2019;12:879–92.

41. Pucker B, Holtgräwe D, Stadermann KB, Frey K, Huettel B, Reinhardt R, et al. A chromosome-level sequence assembly reveals the structure of the Arabidopsis thaliana Nd-1 genome and its gene set. PLOS ONE. 2019;14:e0216233.

42. Slotte T, Hazzouri KM, Ågren JA, Koenig D, Maumus F, Guo Y-L, et al. The Capsella rubella genome and the genomic consequences of rapid mating system evolution. Nat Genet. 2013;45:831–5.

43. Iorizzo M, Ellison S, Senalik D, Zeng P, Satapoomin P, Huang J, et al. A high-quality carrot genome assembly provides new insights into carotenoid accumulation and asterid genome evolution. Nat Genet. 2016;48:657–66.

44. Siadjeu C, Pucker B, Viehöver P, Albach DC, Weisshaar B. High Contiguity de novo Genome Sequence Assembly of Trifoliate Yam (Dioscorea dumetorum) Using Long Read Sequencing. Genes. 2020;11:274.

45. Price MN, Dehal PS, Arkin AP. FastTree 2 – Approximately Maximum-Likelihood Trees for Large Alignments. PLOS ONE. 2010;5:e9490.

46. Katoh K, Standley DM. MAFFT Multiple Sequence Alignment Software Version 7: Improvements in Performance and Usability. Mol Biol Evol. 2013;30:772–80.

47. Pucker B. Automatic identification and annotation of MYB gene family members in plants. BMC Genomics. 2022;23:220.

48. Goodstein DM, Shu S, Howson R, Neupane R, Hayes RD, Fazo J, et al. Phytozome: a comparative platform for green plant genomics. Nucleic Acids Res. 2012;40 Database issue:D1178–86.

49. Caspi R, Billington R, Keseler IM, Kothari A, Krummenacker M, Midford PE, et al. The MetaCyc database of metabolic pathways and enzymes - a 2019 update. Nucleic Acids Res. 2020;48:D445–53.

50. Cheng C-Y, Krishnakumar V, Chan AP, Thibaud-Nissen F, Schobel S, Town CD. Araport11: a complete reannotation of the Arabidopsis thaliana reference genome. Plant J. 2017;89:789–804.

51. Varshney RK, Song C, Saxena RK, Azam S, Yu S, Sharpe AG, et al. Draft genome sequence of chickpea (Cicer arietinum) provides a resource for trait improvement. Nat Biotechnol. 2013;31:240–6.

52. Wu GA, Prochnik S, Jenkins J, Salse J, Hellsten U, Murat F, et al. Sequencing of diverse mandarin, pummelo and orange genomes reveals complex history of admixture during citrus domestication. Nat Biotechnol. 2014;32:656–62.

53. Sato S, Tabata S, Hirakawa H, Asamizu E, Shirasawa K, Isobe S, et al. The tomato genome sequence provides insights into fleshy fruit evolution. Nature. 2012;485:635–41.

54. NCBI. sra-tools. 2020.

55. Pucker B. bpucker/KIPEs: KIPEs v0.35. 2022.

56. Sukumaran J, Holder MT. DendroPy: a Python library for phylogenetic computing. Bioinforma Oxf Engl. 2010;26:1569–71.

57. Gustavsen JA, Pai S, Isserlin R, Demchak B, Pico AR. RCy3: Network biology using Cytoscape from within R. 2019.

58. Letunic I, Bork P. Interactive Tree Of Life (iTOL) v5: an online tool for phylogenetic tree display and annotation. Nucleic Acids Res. 2021;49:W293–6.

59. Xue B. Ensemble Enzyme Prediction Pipeline (E2P2). 2022.

60. Altschul SF, Gish W, Miller W, Myers EW, Lipman DJ. Basic local alignment search tool. J Mol Biol. 1990;215:403–10.

61. Claudel-Renard C, Chevalet C, Faraut T, Kahn D. Enzyme-specific profiles for genome annotation: PRIAM. Nucleic Acids Res. 2003;31:6633–9.

62. Jeske L, Placzek S, Schomburg I, Chang A, Schomburg D. BRENDA in 2019: a European ELIXIR core data resource. Nucleic Acids Res. 2019;47:D542–9.

63. The UniProt Consortium. UniProt: the universal protein knowledgebase in 2021. Nucleic Acids Res. 2021;49:D480–9.

64. Cunningham FX Jr, Pogson B, Sun Z, McDonald KA, DellaPenna D, Gantt E. Functional analysis of the beta and epsilon lycopene cyclase enzymes of Arabidopsis reveals a mechanism for control of cyclic carotenoid formation. Plant Cell. 1996;8:1613–26.

65. Zhao Z, Liu Z, Mao X. Biotechnological Advances in Lycopene β-Cyclases. J Agric Food Chem. 2020;68:11895–907.

66. Lukačin R, Britsch L. Identification of Strictly Conserved Histidine and Arginine Residues as Part of the Active Site in Petunia hybrida Flavanone 3β-Hydroxylase. Eur J Biochem. 1997;249:748–57.

67. Wellmann F, Lukačin R, Moriguchi T, Britsch L, Schiltz E, Matern U. Functional expression and mutational analysis of flavonol synthase from Citrus unshiu. Eur J Biochem. 2002;269:4134–42.

68. Rempel A, Pucker B. BioInfToolServer. BioInfToolServer. 2023. https://pbb-tools.de/. Accessed 23 May 2023.

69. The web framework for perfectionists with deadlines | Django. 2022. https://www.djangoproject.com/. Accessed 1 Jul 2022.

70. Waki T, Takahashi S, Nakayama T. Managing enzyme promiscuity in plant specialized metabolism: A lesson from flavonoid biosynthesis: Mission of a “body double” protein clarified. BioEssays News Rev Mol Cell Dev Biol. 2021;43:e2000164.

71. Wang Z, Wang S, Xu Z, Li M, Chen K, Zhang Y, et al. Highly Promiscuous Flavonoid 3-O-Glycosyltransferase from Scutellaria baicalensis. Org Lett. 2019;21:2241–5.

72. Dewitte G, Walmagh M, Diricks M, Lepak A, Gutmann A, Nidetzky B, et al. Screening of recombinant glycosyltransferases reveals the broad acceptor specificity of stevia UGT-76G1. J Biotechnol. 2016;233:49–55.

73. Ruiz-Sola MÁ, Rodríguez-Concepción M. Carotenoid Biosynthesis in Arabidopsis: A Colorful Pathway. Arab Book. 2012;2012.

74. Just BJ, Santos CAF, Fonseca MEN, Boiteux LS, Oloizia BB, Simon PW. Carotenoid biosynthesis structural genes in carrot (Daucus carota): isolation, sequence-characterization, single nucleotide polymorphism (SNP) markers and genome mapping. Theor Appl Genet. 2007;114:693–704.

75. Dogbo O, Laferriére A, D’Harlingue A, Camara B. Carotenoid biosynthesis: Isolation and characterization of a bifunctional enzyme catalyzing the synthesis of phytoene. Proc Natl Acad Sci U S A. 1988;85:7054–8.

76. Dong H, Deng Y, Mu J, Lu Q, Wang Y, Xu Y, et al. The Arabidopsis Spontaneous Cell Death1 gene, encoding a zeta-carotene desaturase essential for carotenoid biosynthesis, is involved in chloroplast development, photoprotection and retrograde signalling. Cell Res. 2007;17:458–70.

77. Qin G, Gu H, Ma L, Peng Y, Deng XW, Chen Z, et al. Disruption of phytoene desaturase gene results in albino and dwarf phenotypes in Arabidopsis by impairing chlorophyll, carotenoid, and gibberellin biosynthesis. Cell Res. 2007;17:471–82.

78. Bartley GE, Scolnik PA, Beyer P. Two Arabidopsis thaliana carotene desaturases, phytoene desaturase and zeta-carotene desaturase, expressed in Escherichia coli, catalyze a poly-cis pathway to yield pro-lycopene. Eur J Biochem. 1999;259:396–403.

79. Park H, Kreunen SS, Cuttriss AJ, DellaPenna D, Pogson BJ. Identification of the carotenoid isomerase provides insight into carotenoid biosynthesis, prolamellar body formation, and photomorphogenesis. Plant Cell. 2002;14:321–32.

80. Pecker I, Gabbay R, Cunningham FX, Hirschberg J. Cloning and characterization of the cDNA for lycopene beta-cyclase from tomato reveals decrease in its expression during fruit ripening. Plant Mol Biol. 1996;30:807–19.

81. Cunningham FX, Gantt E. One ring or two? Determination of ring number in carotenoids by lycopene epsilon-cyclases. Proc Natl Acad Sci U S A. 2001;98:2905–10.

82. Sun Z, Gantt E, Cunningham FX. Cloning and functional analysis of the beta-carotene hydroxylase of Arabidopsis thaliana. J Biol Chem. 1996;271:24349–52.

83. Tian L, DellaPenna D. Characterization of a second carotenoid beta-hydroxylase gene from Arabidopsis and its relationship to the LUT1 locus. Plant Mol Biol. 2001;47:379–88.

84. Kim J, DellaPenna D. Defining the primary route for lutein synthesis in plants: the role of Arabidopsis carotenoid beta-ring hydroxylase CYP97A3. Proc Natl Acad Sci U S A. 2006;103:3474–9.

85. Kim J-E, Cheng KM, Craft NE, Hamberger B, Douglas CJ. Over-expression of Arabidopsis thaliana carotenoid hydroxylases individually and in combination with a beta-carotene ketolase provides insight into in vivo functions. Phytochemistry. 2010;71:168–78.

86. Tian L, Musetti V, Kim J, Magallanes-Lundback M, DellaPenna D. The Arabidopsis LUT1 locus encodes a member of the cytochrome p450 family that is required for carotenoid epsilon-ring hydroxylation activity. Proc Natl Acad Sci U S A. 2004;101:402–7.

87. Niyogi KK, Grossman AR, Björkman O. Arabidopsis mutants define a central role for the xanthophyll cycle in the regulation of photosynthetic energy conversion. Plant Cell. 1998;10:1121–34.

88. North HM, De Almeida A, Boutin J-P, Frey A, To A, Botran L, et al. The Arabidopsis ABA-deficient mutant aba4 demonstrates that the major route for stress-induced ABA accumulation is via neoxanthin isomers. Plant J Cell Mol Biol. 2007;50:810–24.

89. Ramsay NA, Glover BJ. MYB-bHLH-WD40 protein complex and the evolution of cellular diversity. Trends Plant Sci. 2005;10:63–70.

90. Eddy SR. Accelerated Profile HMM Searches. PLOS Comput Biol. 2011;7:e1002195.

91. Emms DM, Kelly S. SHOOT: phylogenetic gene search and ortholog inference. Genome Biol. 2022;23:85.

