## Additional File 2 for "KIPEs3: Automatic annotation of biosynthesis pathways"

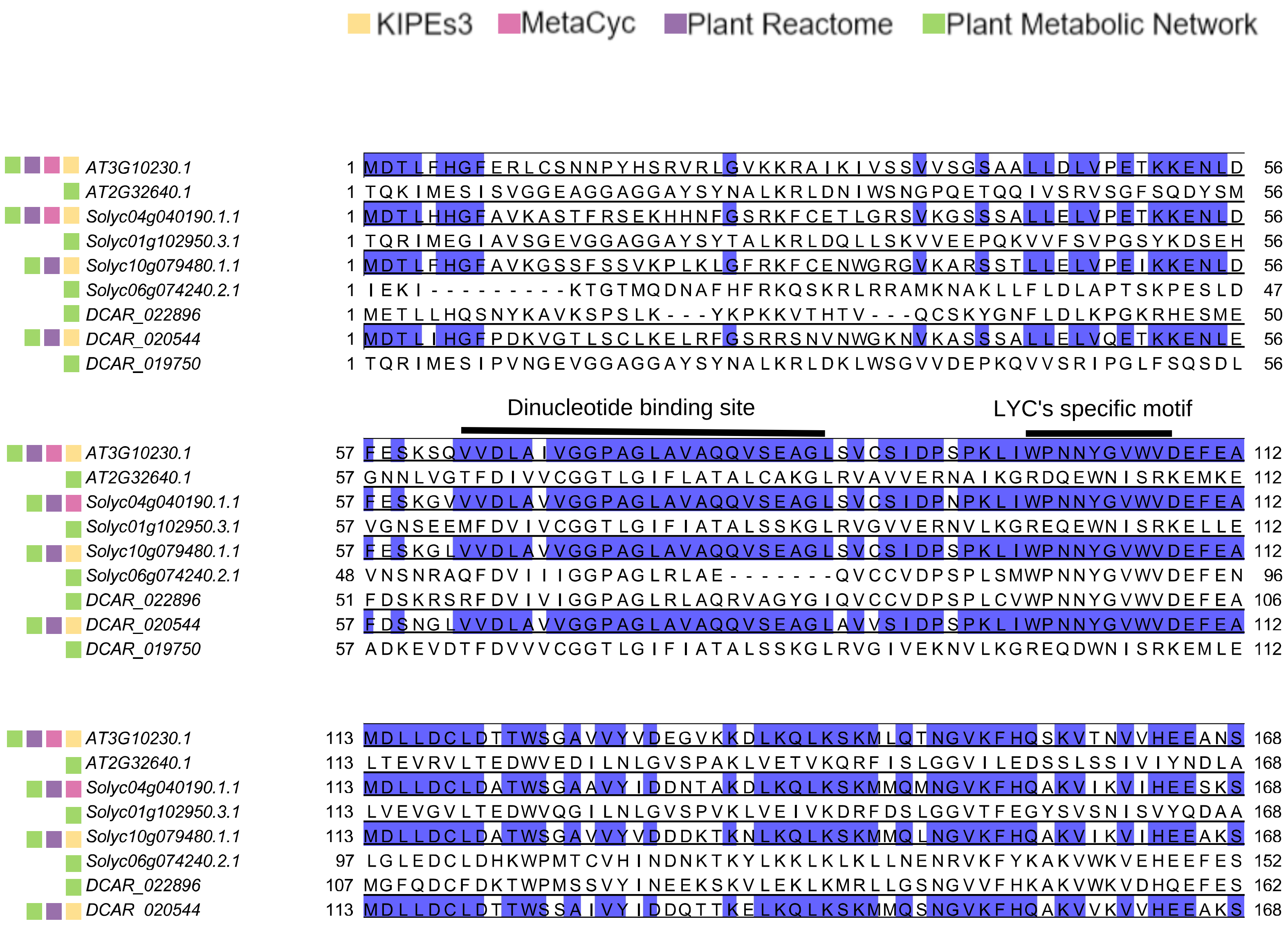
KIPEs3 MetaCyc Plant Reactome Plant Metabolic Network
AT3G10230.1 1 MDTLFHGFERLCSNNPYHSRVRLGVKKRAIKIVSSVVSGSAALLDLVPETKKENLD 56
AT2G32640.1 1 TQKIMESISVGGEAGGAGGAYSYNALKRLDNIWSNGPQETQQIVSRVSGFSQDYSM 56
Solyc04g040190.1.1 1 MDTLHHGFAVKASTFRSEKHHNFGSRKFCETLGRSVKGSSSALLELVPETKKENLD 56
Solyc01g102950.3.1 1 TQRIMEGIAVSGEVGGAGGAYSYTALKRLDQLLSKVVEEPQKVVFSVPGSYKDSEH 56
Solyc10g079480.1.1 1 MDTLFHGFAVKGSSFSSVKPLKLGFRKFCENWGRGVKARSSTLLELVPEIKKENLD 56
Solyc06g074240.2.1 1 IEKI---------KTGTMQDNAFHFRKQSKRLRRAMKNAKLLFLDLAPTSKPESLD 47
DCAR_022896 1 METLLHQSNYKAVKSPSLK---YKPKKVTHTV---QCSKYGNFLDLKPGKRHESME 50
DCAR_020544 1 MDTLIHGFPDKVGTLSCLKELRFGSRRSNVNWGKNVKASSSALLELVQETKKENLE 56
DCAR_019750 1 TQRIMESIPVNGEVGGAGGAYSYNALKRLDKLWSGVVDEPKQVVSRIPGLFSQSDL 56
Dinucleotide binding site
LYC's specific motif
AT3G10230.1 57 FESKSQVVDLAIVGGPAGLAVAQQVSEAGLSVCSIDPSPKLIWPNNYGVWVDEFEA 112
AT2G32640.1 57 GNNLVGTFDIVVCGGTLGIFLATALCAKGLRVAVVERNAIKGRDQEWNISRKEMKE 112
Solyc04g040190.1.1 57 FESKGVVVDLAVVGGPAGLAVAQQVSEAGLSVCSIDPNPKLIWPNNYGVWVDEFEA 112
Solyc01g102950.3.1 57 VGNSEEMFDVIVCGGTLGIFIATALSSKGLRVGVVERNVLKGREQEWNISRKELLE 112
Solyc10g079480.1.1 57 FESKGLVVDLAVVGGPAGLAVAQQVSEAGLSVCSIDPSPKLIWPNNYGVWVDEFEA 112
Solyc06g074240.2.1 48 VNSNRAQFDVIIIGGPAGLRLAE-------QVCCVDPSPLSMWPNNYGVWVDEFEN 96
DCAR_022896 51 FDSKRSRFDVIVIGGPAGLRLAQRVAGYGIQVCCVDPSPLCVWPNNYGVWVDEFEA 106
DCAR_020544 57 FDSNGLVVDLAVVGGPAGLAVAQQVSEAGLAVVSIDPSPKLIWPNNYGVWVDEFEA 112
DCAR_019750 57 ADKEVDTFDVVVCGGTLGIFIATALSSKGLRVGIVEKNVLKGREQDWNISRKEMLE 112
AT3G10230.1 113 MDLLDCLDTTWSGAVVYVDEGVKKDLKQLKSKMLQTNGVKFHQSKVTNVVHEEANS 168
AT2G32640.1 113 LTEVRVLTEDWVEDILNLGVSPAKLVETVKQRFISLGGVILEDSSLSSIVIYNDLA 168
Solyc04g040190.1.1 113 MDLLDCLDATWSGAAVYIDDNTAKDLKQLKSKMMQMNGVKFHQAKVIKVIHEESKS 168
Solyc01g102950.3.1 113 LVEVGVLTEDWVQGILNLGVSPVKLVEIVKDRFDSLGGVTFEGYSVSNISVYQDAA 168
Solyc10g079480.1.1 113 MDLLDCLDATWSGAVVYVDDDKTKNLKQLKSKMMQLNGVKFHQAKVIKVIHEEAKS 168
Solyc06g074240.2.1 97 LGLEDCLDHKWPMTCVHINDNKTKYLKKLKLKLLNENRVKFYKAKVWKVEHEEFES 152
DCAR_022896 107 MGFQDCFDKTWPMSSVYINEEKSKVLEKLKMRLLGSNGVVFHKAKVWKVDHQEFES 162
DCAR_020544 113 MDLLDCLDTTWSSAIVYIDDQTTKELKQLKSKMMQSNGVKFHQAKVVKVVHEEAKS 168

```
DCAR_019750         113 LVEVGILEEEWVENILNLGVSPSKLIEKMKTRFNSFDGVILEGLGVSSICVYDDAA 168

AT3G10230.1         169 TVVCSDGVKIQASVVLDATGFSRCLVQYDKPYNPGYQVAYGIVAHPFDVDKMRDKH 224
AT2G32640.1         169 VMQLSKGDTLSSRLVIDAMGFSPILKQIKRGRKPGMCLVVGSCAHGFKENSSSDVI 224
Solyc04g040190.1.1  169 MLICNDGITIQATVVLDATGFSRSLVQYDKPYNPGYQVAYGILAHPFDVNKMRDSH 224
Solyc01g102950.3.1  169 VLQLKEGKTLFSRLIIDAMGFSPIVKQIRCGRKPGMCLVVGTCCRGFKENSTSDVI 224
Solyc10g079480.1.1  169 MLICSDGVTIQATVVLDATGFSRCLVQYDKPYNPGYQVAYGILAHPFDTSKMRDSH 224
Solyc06g074240.2.1  153 SIVCDDGKKIRGSLVVDASGFASDFIEYDRPRNHGYQIAHGVLVHPFDLDKMRDSH 208
DCAR_022896         163 SILCDDGKEFKASLIVDASGFASTFVDYDKPRNHGYQLAHGILAHPFELDRMRDSH 218
DCAR_020544         169 LLICNDGVTIQAAVVLDATGFSRCLVQYDKPYNPGYQVAYGIVAHPFDVNKMRDSH 224
DCAR_019750         169 ILQLDSGKRLSSRLVIDAMGFSPVVKQIRGGRKPGFCLVVGSCCRGFKDNKTSDVI 224

                                  LYC-b CAD
                             CMI              CMI
AT3G10230.1         225 LDSYPELKERNSKIPTFLYAMPFSSNRIFLEETSLVARPGLRMEDIQERMAARLKH 280
AT2G32640.1         225 YSSSSVTRVADSNVQLFWEAFPAGSGPLFTYTEPQSTSPSLDLLEEYWKLMPKYQG 280
Solyc04g040190.1.1  225 LKNNTDLKERNSRIPTFLYAMPFSSNRIFLEETSLVARPGLRIDDIQERMVARLNH 280
Solyc01g102950.3.1  225 FSSATAKEVGQSLVQYFWEAFPAGSGPIFTYVDPQPGSPQLELLEDYWDLMPKYQG 280
Solyc10g079480.1.1  225 LNNNVKLKERNRKVPTFLYAMPFSSNRIFLEETSLVARPGLRMDDIQERMVARLSH 280
Solyc06g074240.2.1  209 LGNEPYLRVNNAKEPTFLYAMPFDRDLVFLEETSLVSRPVLSYMEVKRRMVARLRH 264
DCAR_022896         219 LGNEPALRFANAKSPTFLYAMPFDSNLIFLEETSLVSRPALSYKEVKLRMAARLRH 274
DCAR_020544         225 LNGNTELKERNSKIPTFLYAMPFSSDRIFLEETSLVARPGLAMGDIQERMVARLRH 280
DCAR_019750         225 YSSAEVMQVGESQVQYFWEAFPAGSGLMFTYVDPQPGSPKLELLEDYWNLMPDYQG 280

                                                          LYC-b CAD
                             LYC-b CAD              CMII
AT3G10230.1         281 LGINVKRIEEDERCVIPMGGPLPVLPQRVVGIGGTAGMVHPSTGYMVARTLAAAPI 336
AT2G32640.1         281 VSLDELEILRVVYGIFPTNSPLPAAFDRVLQFGDASGIQSPVSGSLTRHLGRLSNG 336
Solyc04g040190.1.1  281 LGIKVKSIEEDEHCLIPMGGPLPVLPQRVVGIGGTAGMVHPSTGYMVARTLAAAPV 336
Solyc01g102950.3.1  281 VSFDDLEILRIIFGIFPTDSPLPAAFDRILQFGDASGIQSPVSGSLTRHLGRLTTG 336
Solyc10g079480.1.1  281 LGIKVTSIEEDEQCVIPMGGPLPVIPQRVVGIGGTAGMVHPSTGYMVARTLAAAPV 336
```

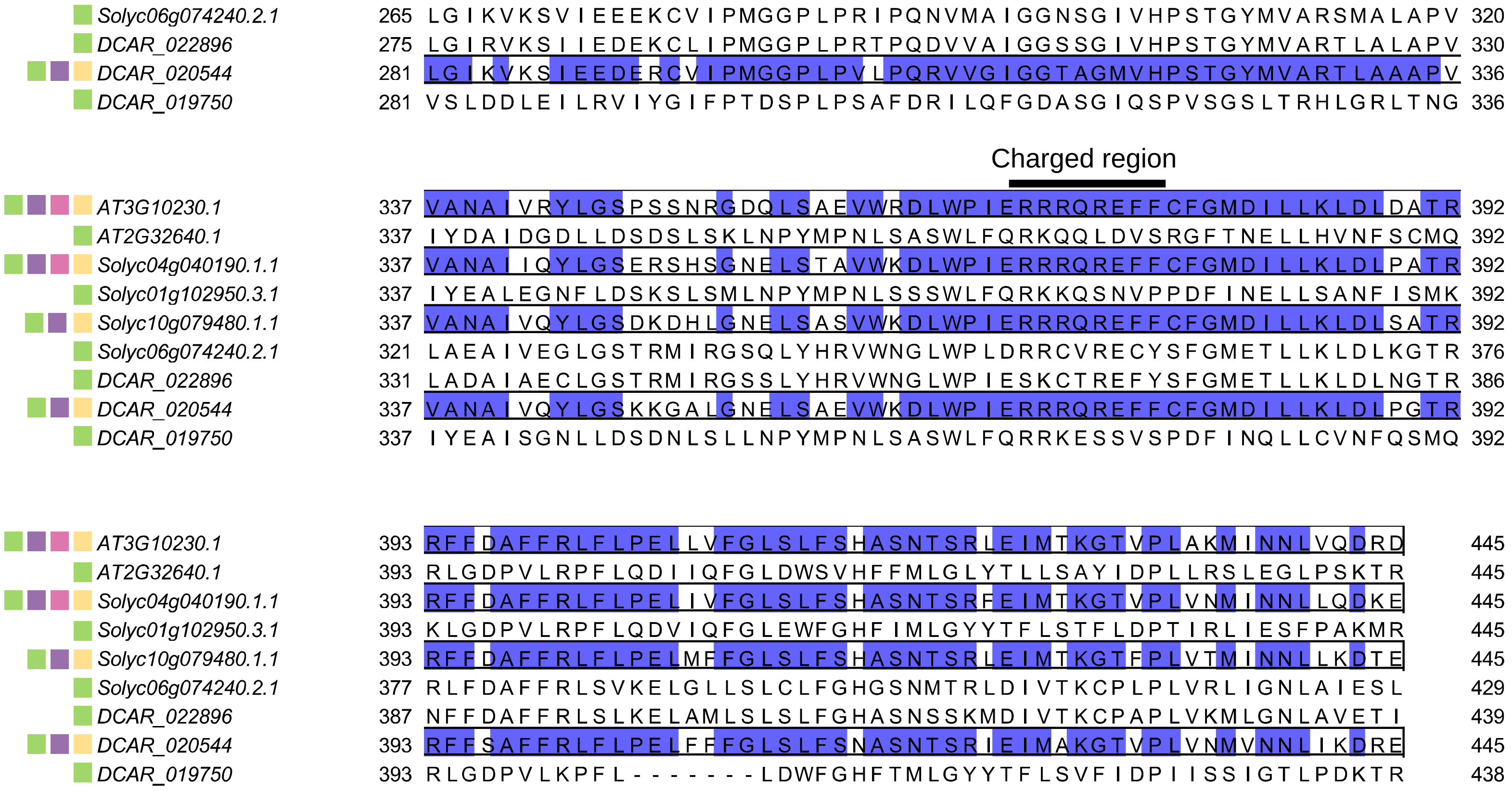
Solyc06g074240.2.1 265 LGIKVKSVIEEEKCVIPMGGPLPRIPQNVMAIGGNSGIVHPSTGYMVARSMALAPV 320
DCAR_022896 275 LGIRVKSIIEDEKCLIPMGGPLPRTPQDVVAIGGSSGIVHPSTGYMVARTLALAPV 330
DCAR_020544 281 LGIKVKSIEEDERCVIPMGGPLPVLPQRVVGIGGTAGMVHPSTGYMVARTLAAAPV 336
DCAR_019750 281 VSLDDLEILRVIYGIFPTDSPLPSAFDRILQFGDASGIQSPVSGSLTRHLGRLTNG 336
Charged region
AT3G10230.1 337 VANAIVRYLGSPSSNRGDQLSAEVWRDLWPIERRRQREFFCFGMDILLKLDLDATR 392
AT2G32640.1 337 IYDAIDGDLLDSDSLSKLNPYMPNLSASWLFQRKQQLDVSRGFTNELLHVNFSCMQ 392
Solyc04g040190.1.1 337 VANAIIQYLGSERSHSGNELSTAVWKDLWPIERRRQREFFCFGMDILLKLDLPATR 392
Solyc01g102950.3.1 337 IYEALEGNFLDSKSLSMLNPYMPNLSSSWLFQRKKQSNVPPDFINELLSANFISMK 392
Solyc10g079480.1.1 337 VANAIVQYLGSDKDHLGNELSASVWKDLWPIERRRQREFFCFGMDILLKLDLSATR 392
Solyc06g074240.2.1 321 LAEAIVEGLGSTRMIRGSQLYHRVWNGLWPLDRRCVRECYSFGMETLLKLDLKGTR 376
DCAR_022896 331 LADAIAECLGSTRMIRGSSLYHRVWNGLWPIESKCTREFYSFGMETLLKLDLNGTR 386
DCAR_020544 337 VANAIVQYLGSKKGALGNELSAEVWKDLWPIERRRQREFFCFGMDILLKLDLPGTR 392
DCAR_019750 337 IYEAISGNLLDSDNLSLLNPYMPNLSASWLFQRRKESSVSPDFINQLLCVNFQSMQ 392
AT3G10230.1 393 RFFDAFFRLFLPELLVFGLSLFSHASNTSRLEIMTKGTVPLAKMINNLVQDRD 445
AT2G32640.1 393 RLGDPVLRPFLQDIIQFGLDWSVHFFMLGLYTLLSAYIDPLLRSLEGLPSKTR 445
Solyc04g040190.1.1 393 RFFDAFFRLFLPELIVFGLSLFSHASNTSRFEIMTKGTVPLVNMINNLLQDKE 445
Solyc01g102950.3.1 393 KLGDPVLRPFLQDVIQFGLEWFGHFIMLGYYTFLSTFLDPTIRLIESFPAKMR 445
Solyc10g079480.1.1 393 RFFDAFFRLFLPELMFFGLSLFSHASNTSRLEIMTKGTFPLVTMINNLLKDTE 445
Solyc06g074240.2.1 377 RLFDAFFRLSVKELGLLSLCLFGHGSNMTRLDIVTKCPLPLVRLIGNLAIESL 429
DCAR_022896 387 NFFDAFFRLSLKELAMLSLSLFGHASNSSKMDIVTKCPAPLVKMLGNLAVETI 439
DCAR_020544 393 RFFSAFFRLFLPELFFFGLSLFSNASNTSRIEIMAKGTVPLVNMVNNLIKDRE 445
DCAR_019750 393 RLGDPVLKPFL--------LDWFGHFTMLGYYTFLSVFIDPIISSIGTLPDKTR 438
