## Supplementary figures and images for "KIPEs3: Automatic annotation of biosynthesis pathways"

### Additional File 3

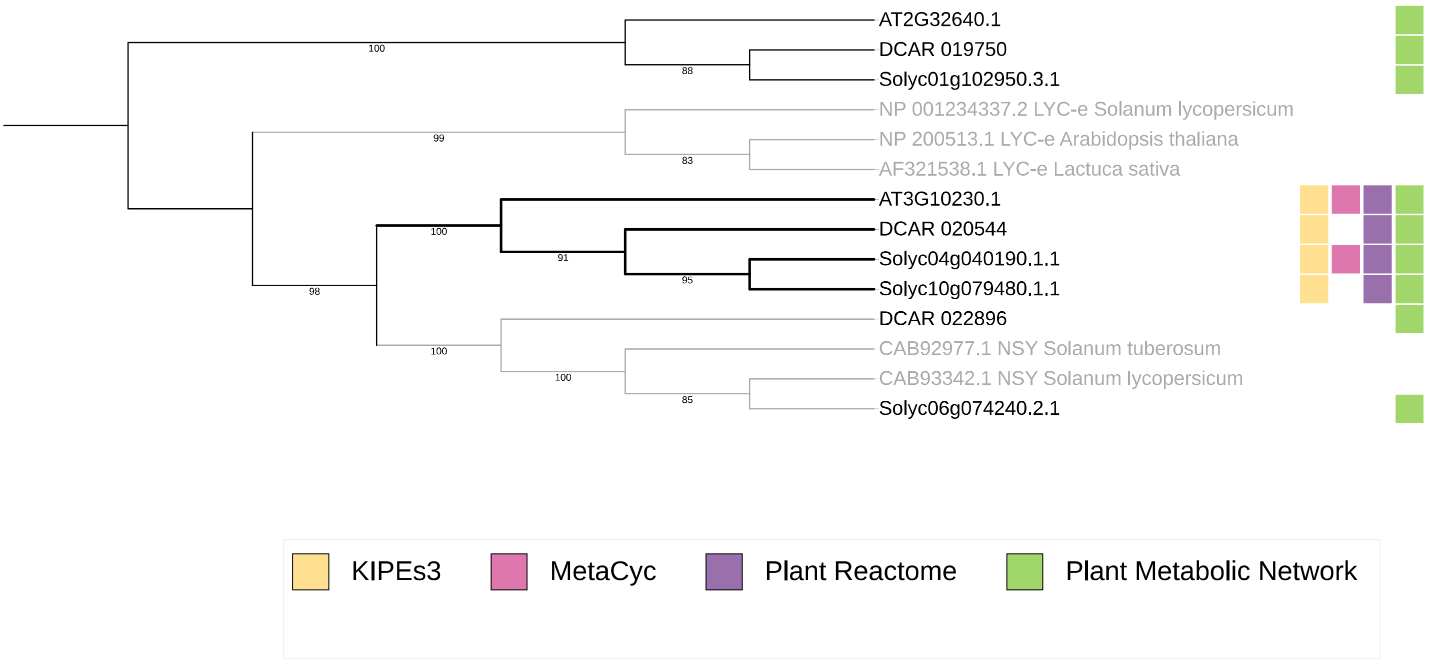

### Additional File 4

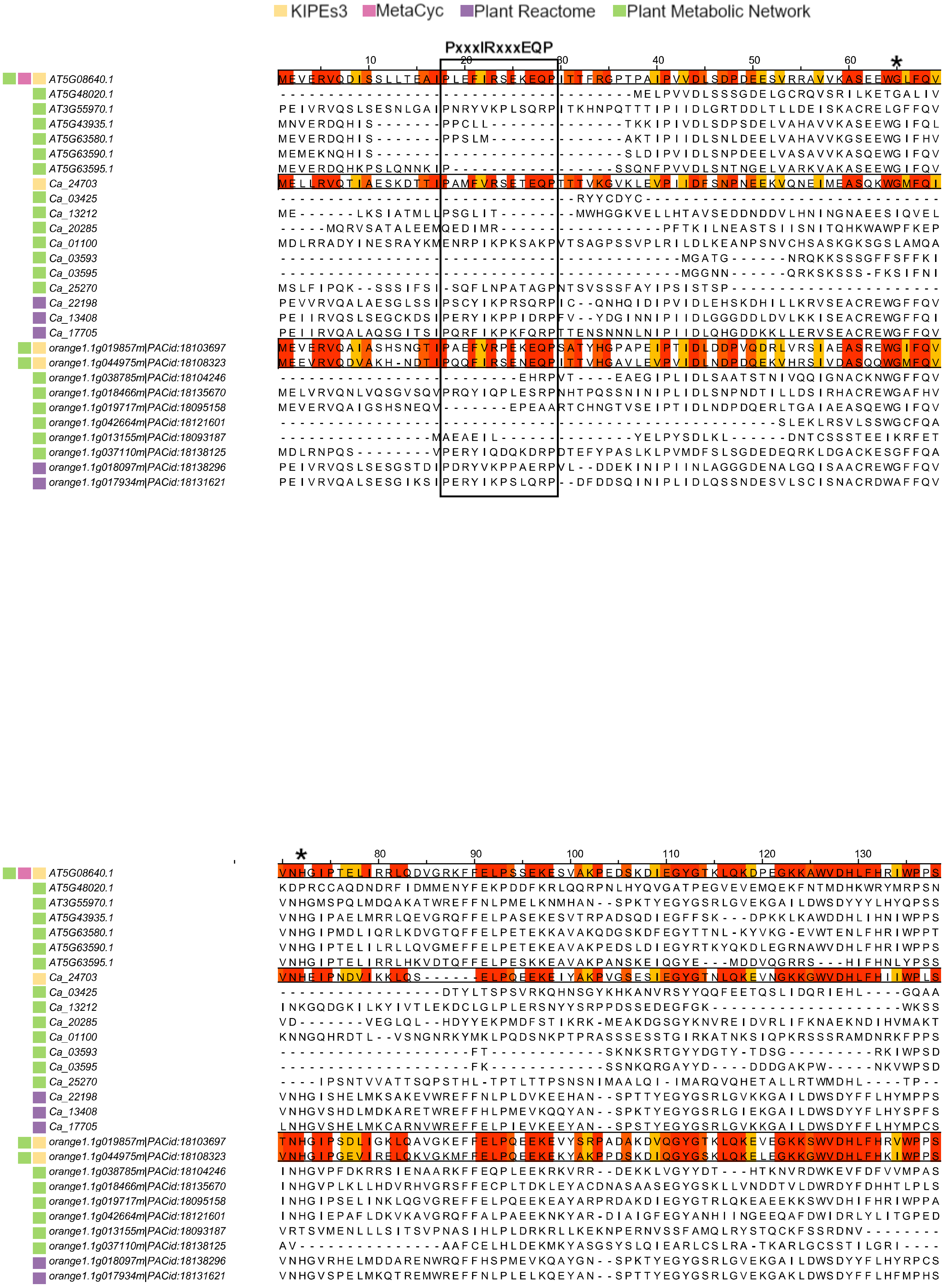

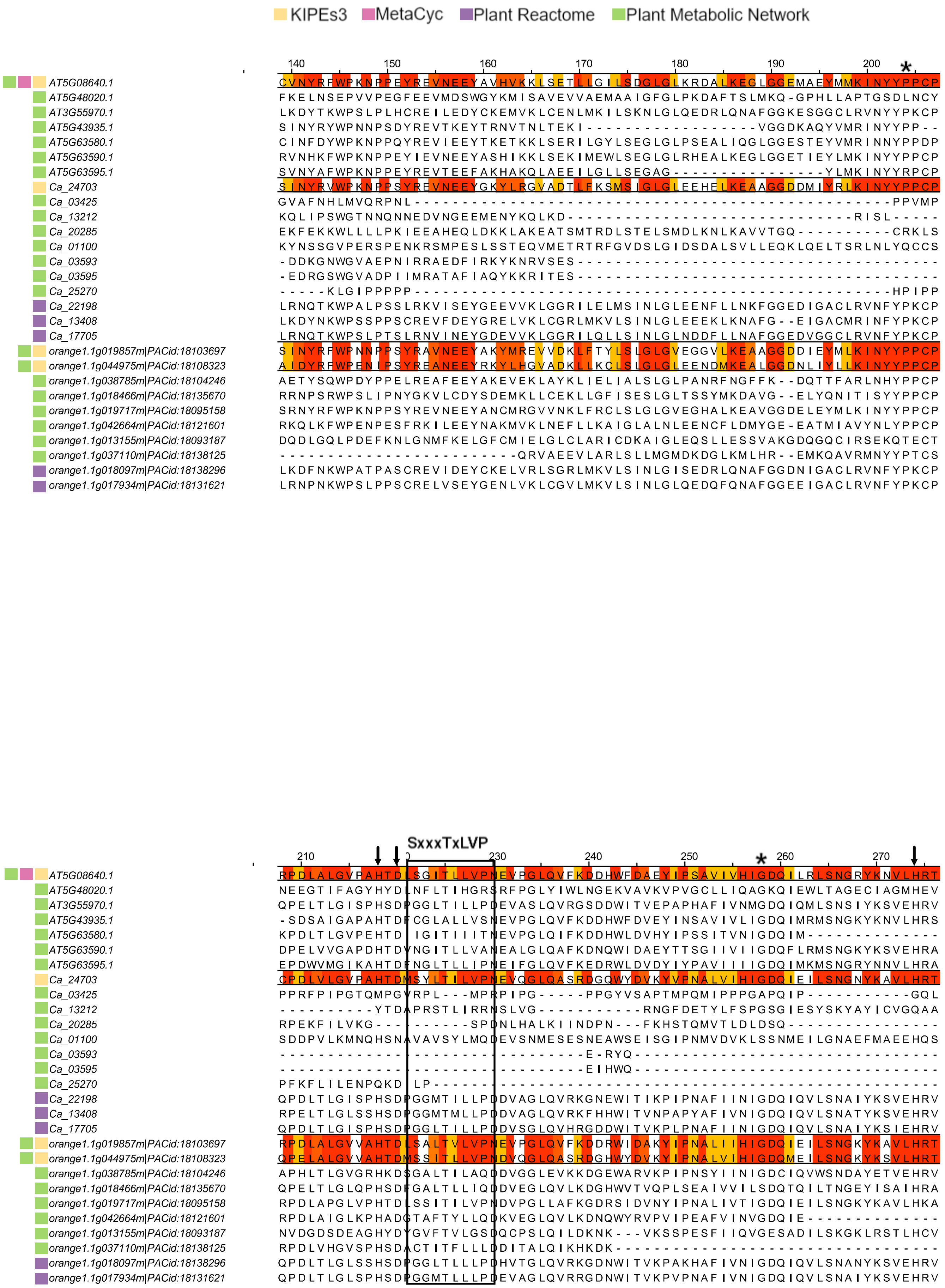

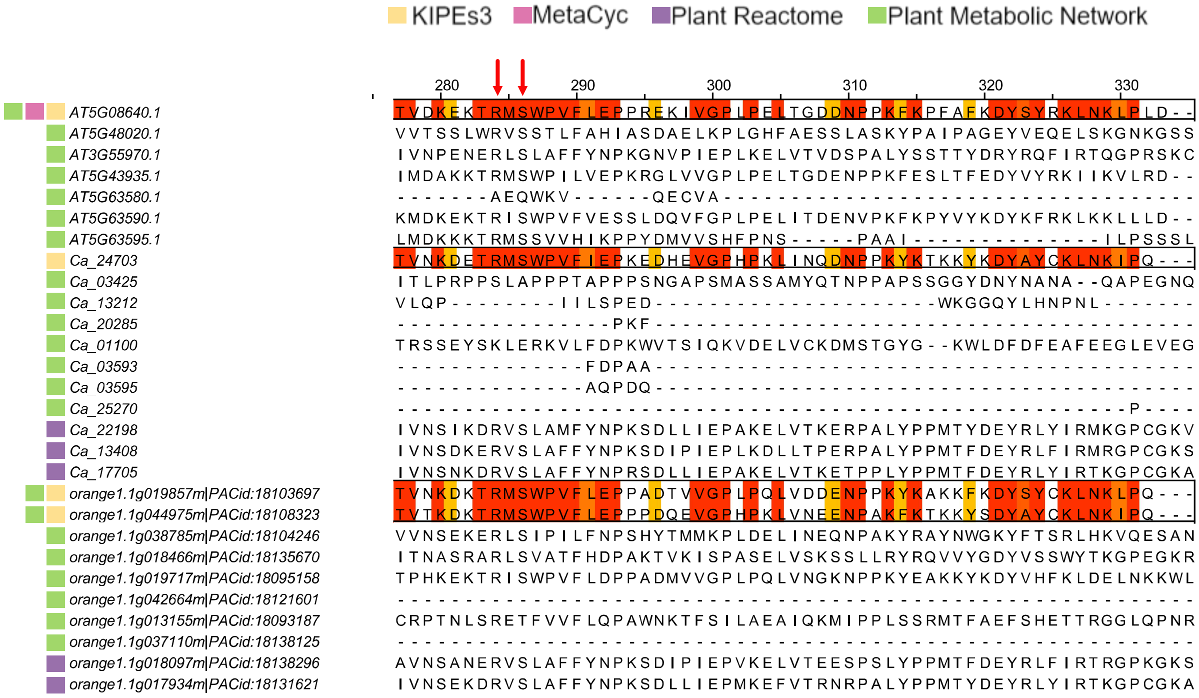
